# Dopamine D1 receptor expression in prefrontal parvalbumin neurons influences distractibility across species

**DOI:** 10.1101/2024.06.15.599163

**Authors:** MKP Joyce, TG Ivanov, FM Krienen, JF Mitchell, S Ma, W Inoue, AS Nandy, D Datta, A Duque, J Arellano, R Gupta, G Gonzalez-Burgos, DA Lewis, N Sestan, SA McCarroll, J Martinez-Trujillo, S Froudist-Walsh, AFT Arnsten

## Abstract

Marmosets and macaques are common non-human primate models of cognition, yet marmosets appear more distractible and perform worse in cognitive tasks. The dorsolateral prefrontal cortex (dlPFC) is pivotal for sustained attention, and prior macaque research suggests that dopaminergic modulation and inhibitory parvalbumin (PV) neurons could contribute to distractibility. Thus, we compared the two species using a visual fixation task with distractors, performed molecular and anatomical analyses in dlPFC, and linked functional microcircuitry with cognitive performance using computational modeling. We found that marmosets are more distractible than macaques, and that marmoset dlPFC PV neurons contain higher levels of dopamine-1 receptor (D1R) transcripts and protein, similar to their levels in mice. The modeling indicated that higher D1R expression in marmoset dlPFC PV neurons may increase distractibility by making dlPFC microcircuits more vulnerable to disruptions of their task-related persistent activity, especially when dopamine is released in dlPFC in response to unexpected salient stimuli.

**Declaration of Interests:** The authors have nothing to declare.

**Author Contributions:** AFTA, MKPJ, TGI, and SFW designed the study; MKPJ collected and analyzed anatomical data; TGI designed the computational framework with guidance from RG and SFW; TGI performed the modeling simulations and data analysis; FM and SAM collected and analyzed transcriptomics data; GGB and DAL provided macaque brain tissue for transcriptomics performed by FM; SM and NS collected and analyzed phylogenetic transcriptomics dataset; JFM and ASN performed and analyzed the behavioral testing; IW and JMT provided the marmoset tissue for immunofluorescence; MKPJ, TGI, SFW, and AFTA wrote the first draft and all authors revised and edited subsequent drafts of the article.

## Introduction

Marmosets are an increasingly attractive non-human primate model due to their small size, ease of handling, short breeding cycle, brief lifespan, and amenability to emerging genetic and molecular technologies (e.g.,^**1**^). The use of marmosets has already proven successful in modeling many psychiatric and mental processes^**2-4**^. Yet the extent of their translational validity and similarity to macaques in cognitive domains is still under investigation, as very few studies have directly compared the two species in the same tasks^5, 6^. One early study^**7**^ found that marmosets required more trials to match macaque performance in a delayed-response working memory task, and marmoset performance, compared to that of macaques, worsened more with longer delays. Additionally, marmosets invariably failed the task if they became distracted away from the screen, while macaques could overcome environmental distractions and respond correctly^**7**^. More recently, assays of attentional regulation showed that macaques^**8**^ have a superior ability to regulate attention shifting compared to marmosets^**9**^. These data are consistent with anecdotal reports that high levels of distractibility in marmosets can interfere with training^**10**^. Although a more comprehensive comparison of cognitive abilities across these species remains necessary, the evidence suggests that marmosets perform more poorly in tasks with higher cognitive demands, and that they may be more susceptible than macaques to distractibility. These cross-species differences in cognitive function necessitate a comparison of the neural circuitry supporting these processes in marmosets and macaques.

Core cognitive processes, such as working memory maintenance and sustained attention, may rely on only partially overlapping neuronal populations in layer III dlPFC^**11**, **12**^, and recent findings suggest a strong dissociation^**13**^. Nevertheless, these processes may have a common neurophysiological basis in the persistent firing activity of layer III pyramidal neurons. Specifically, studies of macaque dlPFC have revealed that extensive recurrent excitation among pyramidal neurons in specialized microcircuits of layer III dlPFC sustains persistent activity across a working memory delay period while subjects visually fixate on a central point, and even in the presence of distractors^**14**, **15**^, as well as during visual fixation alone without additional working memory demands^**16-21**^. These layer III microcircuits are considered essential for actively maintaining covert mental representations of target stimuli, even in the absence of external input, as well as task rules—e.g., to fixate overtly on a central point. Inhibitory, GABAergic PV neurons have also been central to early theories of the cellular basis of working memory maintenance and sustained attention^**14**, **15**, **22**^, although their exact contributions to cognitive function and particularly distractibility are still unclear. Evidence suggests that fast-spiking, putative PV neurons may be actively recruited during the delay period to sculpt the spatial tuning of excitatory population activity^**14**, **23**, **24**^, and adequate spatial tuning is critical for distractor resistance. PV neurons likely provide feedback inhibition onto the same pyramidal neurons that excite them^**25-33**^. They may also provide feedforward lateral inhibition onto pyramidal neurons with different spatial tuning^**25-33**^, which could include pyramidal neurons with selectivities that are either similar to or dissimilar from the selectivity of the pyramidal neurons that excite the given PV neuron. Whether the microcircuit components contributing to working memory maintenance and sustained attention in macaque dlPFC are present in marmoset dlPFC remains largely unknown. Marmosets have a smaller dlPFC than macaques^**34-36**^, and marmoset prefrontal pyramidal neurons have fewer spines on the basal dendrites^**37**^, which could be necessary for the recurrent excitation that supports persistent activity. Nevertheless, dlPFC delay-related persistent activity was recently verified in marmosets^**38**^, suggesting that they have the key microcircuit components to support working memory maintenance, which may also extend to sustained attention.

Neuromodulators such as dopamine (DA) have powerful effects on dlPFC function in macaques^**39**^ and marmosets^**40**^. DA is released in the dlPFC to salient stimuli, whether they are rewarding or aversive^**41**^. In macaques, DA largely influences dlPFC function via D1Rs^**39**, **42**^, with an inverted-U dose-response, where mid-range levels of D1R stimulation facilitate working memory maintenance and sustained attention^**43**, **44**^, while higher levels impair them^**45**, **46**^. In macaque dlPFC, D1Rs predominate on the dendritic spines of layer III pyramidal neurons^**42**, **44**, **47**^, but they are also found on the dendrites of PV neurons^**48**^. *Ex vivo* physiological studies in rodents^**49**, **50**^and macaques^**51**^ have shown that D1R stimulation increases the intrinsic excitability of fast-spiking, putative PV neurons, which could affect their roles in the structured inhibition and spatial tuning of excitatory pyramidal neurons *in vivo*. A recent study using iontophoresis and electrophysiology in macaque dlPFC showed that DA, via D1Rs, modulates fast-spiking, likely PV, neurons during delay periods with distractor challenges^**52**^. Given the D1R-mediated increase in the excitability of fast-spiking, putative PV neurons, we hypothesized that higher D1R expression levels in layer III dlPFC PV neurons in marmosets, compared to macaques, may significantly degrade distractor resistance if there is excessive inhibition of pyramidal neuron firing. Because these layer III microcircuits are altered in schizophrenia^**53**^, stage III/IV Alzheimer’s disease^**54**^, and stress and mood disorders^**55**^, understanding their common and divergent properties across primates is critical when modeling neural mechanisms of cognitive disorders.

Here, we first compared the ability of marmosets and macaques to maintain gaze on a central point in the presence of visual distractors. Next, we assessed D1R expression in layer III dlPFC PV neurons across the two species by mining and integrating data from existing single nucleus RNA sequencing (snRNA-seq) datasets, including data from mice and humans, and then performing immunohistochemistry and quantitative image analysis. Finally, we incorporated the anatomical measurements of D1R expression into computational models of covert visuospatial working memory maintenance and overt sustained attention during visual fixation to simulate the effect of higher D1R expression in inhibitory fast-spiking, putative PV neurons in layer III dlPFC microcircuits. Our results demonstrate that marmosets exhibit more distractibility than macaques in a visual fixation task, and that marmosets have higher D1R expression in layer III dlPFC PV neurons. Lastly, our modeling suggests that the higher D1R expression in layer III dlPFC PV neurons may contribute to higher marmoset distractibility by causing more prevalent disruptions of task-related persistent activity, particularly when DA is released in dlPFC in response to unexpected salient stimuli. These disruptions would, in turn, result in reflexive behavioral responses. Our results suggest fundamental species differences in the effect of DA on dlPFC microcircuits and offer insights into neurophysiological basis of distractibility in working memory maintenance and sustained attention.

## Results

### Marmosets are more distractible than macaques in a visual fixation task

To quantify differences in distractibility between marmosets and macaques, we used a visual fixation task (Figure 1A-B) requiring overt sustained attention and distractor resistance, but no working memory maintenance. Specifically, we examined the ability of subjects to hold visual fixation on a central point while spatially unpredictable, distracting stimuli flashed in the visual periphery. This task is commonly used with rhesus macaques to map receptive field properties of neurons in visual cortical areas^**56**^, and marmosets can perform it with a reasonable degree of success^**57**^. We analyzed data previously collected from marmosets and macaques performing this task under comparable visual stimulation conditions^**56**, **57**^. Animals were required to maintain fixation within a 1-2° radius around the fixation point while high contrast Gabor stimuli flashed at random positions in the periphery at 60 Hz. Monkeys were rewarded with a drop of juice for holding fixation at increasingly longer periods up to a maximum of 3 seconds.

**Figure 1.**
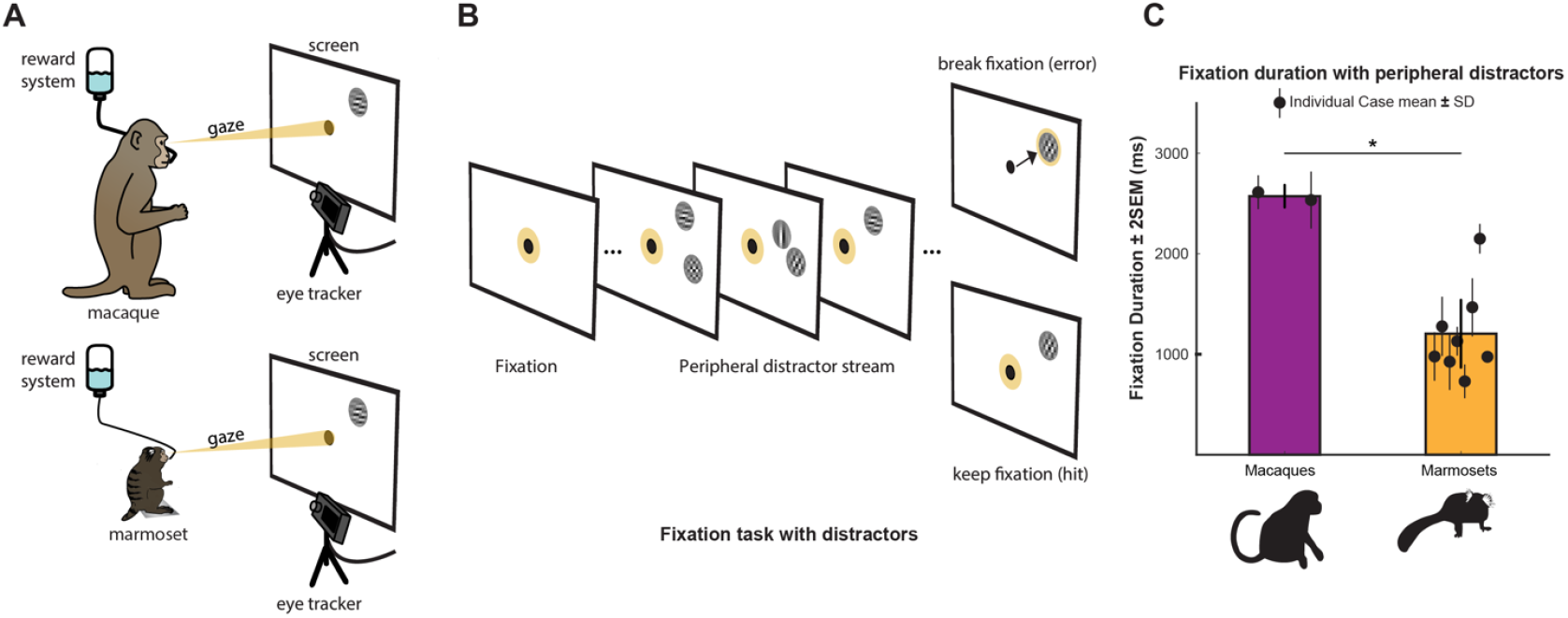
Macaques held central fixation in the presence of peripheral distractors more than twice as long as marmosets. ***A***, Schematic of the task setup. A macaque (top) and a marmoset (bottom) face a screen, and an eye tracker monitors their eye movements. Monkeys receive a juice reward for maintaining fixation at the central point in the presence of peripheral distractors. ***B***, Schematic of the task progression. A point appears at the center of the screen, and the monkey fixes its gaze (yellow) on the central point. Gabor stimuli flash in and out of the periphery, and the monkey is rewarded for maintaining its fixation on the central target (“hit”) despite the distracting peripheral stimuli. ***C***, Time held fixating at a central point in the presence of visual distractors across species. Bars indicate group mean fixation duration for two macaques (purple) and for eight marmosets (orange). Thick black lines depict ± 2 standard error of the mean (SEM); Black circles and thin lines depict individual subject mean fixation duration ± standard deviation (SD). ^*^, p < 0.05, two-tailed Ranksum test.

The macaques could ignore the distractors for over twice as long as any of the marmosets (Figure 1C), holding fixation for a median duration of 2.57 seconds as compared to 1.05 seconds in marmosets, which was a significant cross-species difference (two-tailed Ranksum test, p = 0.044). Although individual marmoset performance varied considerably, no single marmoset achieved performance equivalent to either of the macaques (individuals shown with black circles, Figure 1C). These data suggest that even when the cross-species comparison task has no working memory maintenance demands, a fundamental difference in distractibility is detectable between the species.

### Marmosets have higher DRD1 expression in dlPFC PVALB neurons than macaques

Analyses from independent snRNA-seq datasets revealed that *DRD1* (the gene encoding D1R) expression in *PVALB* (the gene encoding PV) neuron clusters of the marmoset dlPFC is markedly higher than in macaque dlPFC (Figure 2). First, we compared *DRD1* expression in *PVALB* neuron clusters of young adult marmosets^**58**^ and macaques^**59**^, and found higher expression of *DRD1* in *PVALB* clusters from marmoset compared to macaque (Figure 2A). Less than 1% of macaque *PVALB* nuclei were *DRD1*+, while about 20% of the marmoset *PVALB* nuclei were *DRD1*+. Then, our second dataset^**60**, **61**^ corroborated and extended these findings, demonstrating that *DRD1* expression in *PVALB* clusters of marmosets exceeded expression in macaques, and that marmoset dlPFC and mouse frontal cortex exhibit high *DRD1* expression in *PVALB* clusters, (Figure 2B), whereas human dlPFC had the lowest expression. Overall, these results suggest an inverse relationship between *DRD1* expression and phylogenetic position among the species studied here.

**Figure 2.**
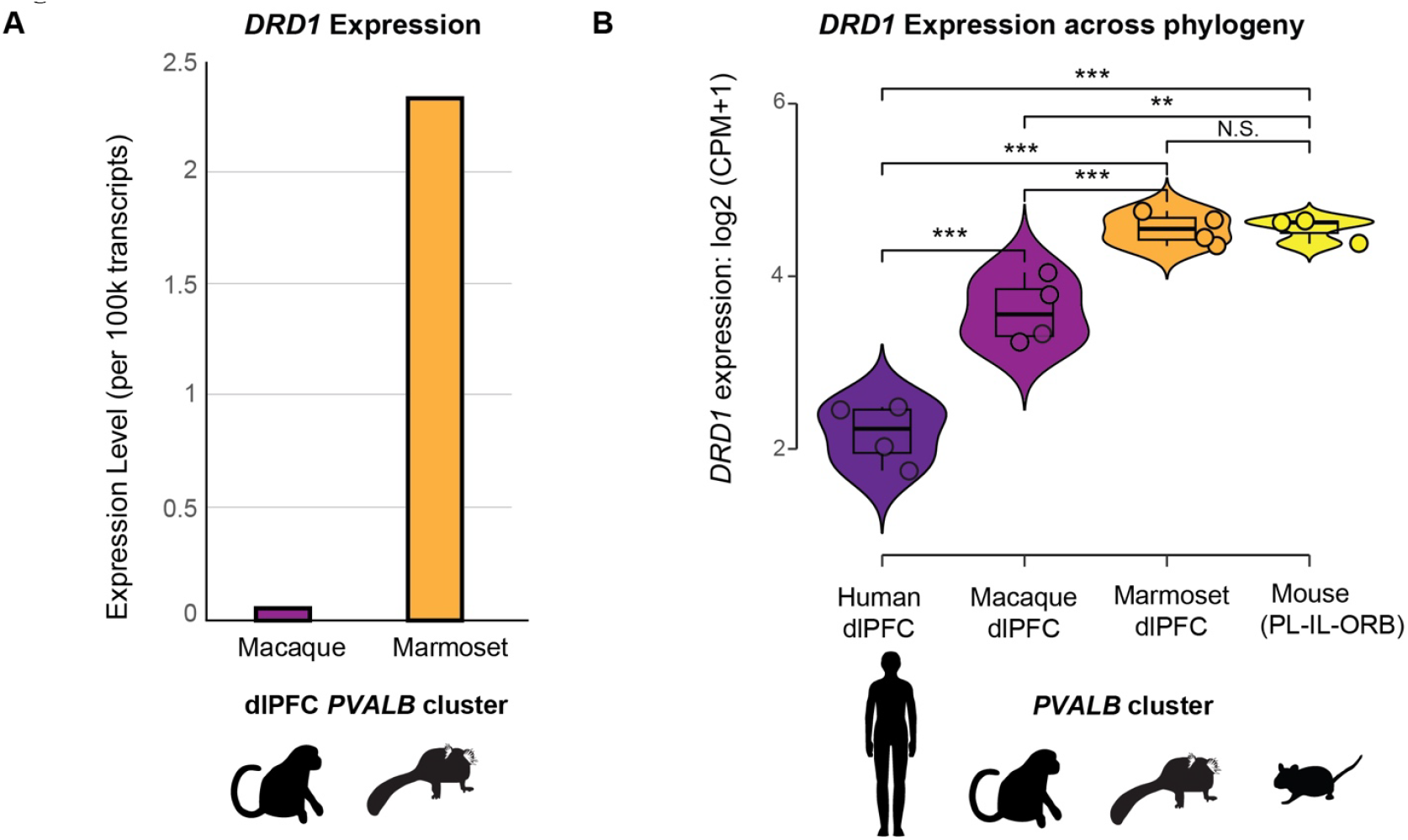
Marmoset dlPFC *PVALB* neurons have higher *DRD1* expression than macaque *PVALB* neurons. ***A***, *DRD1* expression in *PVALB* clusters of the dlPFC in macaques^**59**^ (purple) and marmosets^**58**^ (orange), depicted as expression per 100,000 transcripts. ***B***, Library size-normalized expression of *DRD1* in adult cortical *PVALB* neurons across human dlPFC (dark purple), macaque dlPFC (plum), marmoset dlPFC (orange), and mouse frontal cortex (yellow). Data mined from Ma *et al*.^**61**^ and Yao *et al*.^**60**^. Due to the lack of dlPFC homology in mice, we included mouse PL-IL-ORB (prelimbic, infralimbic and orbital regions, yellow) regions, which are analogous substrates for some primate dlPFC cognitive functions. Each dot represents a pseudobulk donor and the significance of the gene expression differences was tested by edgeR^**134**^. CPM, counts per million; ^***^, p < 0.001; ^**^, p < 0.01; N.S., not significant.

### Marmosets have higher D1R expression in layer III dlPFC PV neurons than macaques

To examine the *in situ* expression of D1R protein in layer III dlPFC, we labeled D1R and PV using immunohistochemistry (Figure 3A-B). In both species, D1R labeling was observed in the cytoplasmic and nuclear regions of PV neurons and putative pyramidal neurons, consistent with previous findings^**48**, **62**, **63**^.

**Figure 3.**
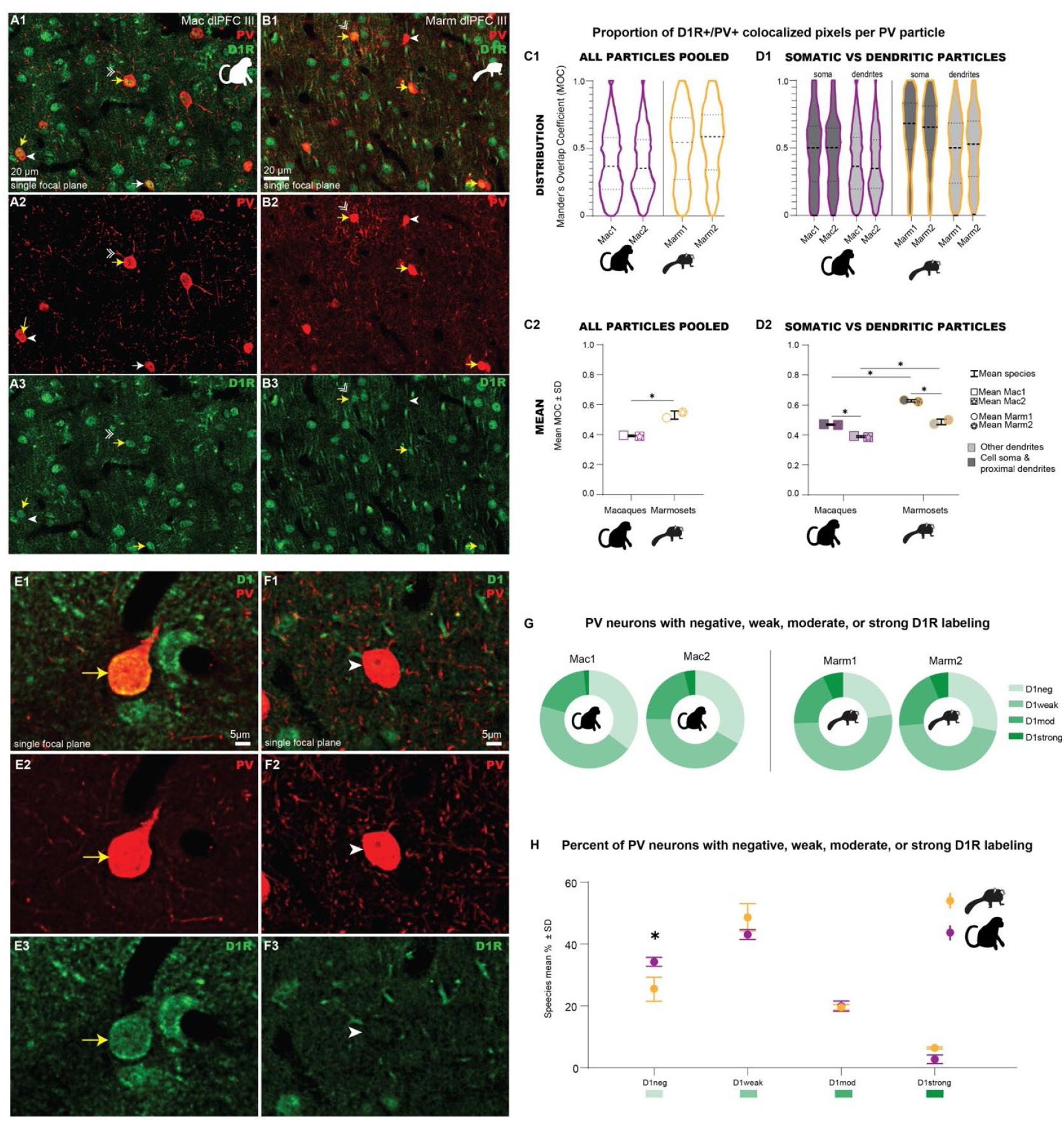
Marmosets have higher correlates of D1R protein expression in layer III dlPFC PV neurons than macaques. ***A-B***, Confocal photomicrographs with brightness and contrast adjustments depicting a single focal plane imaged in layer III of the macaque **(A)** and marmoset **(B)** dlPFC labeled for PV (red) and D1R (green). Images depict some PV neurons robustly expressing D1R (yellow arrows), or not expressing D1R (white arrowhead). Double-headed white arrow heads depict proximal PV dendrites, defined by being traceable to a parent PV soma. ***C1***, Violin plots with medians and quartiles, depicting kernel density of the MOC pooled across all PV particles by subject. MOC roughly measures the proportion of D1R+/PV+ pixels in each PV particle. Macaques, purple; marmosets, orange. ***C2***, Species MOC means for all PV particles (marmosets MOC 0.53±0.03, macaques 0.39±0.01; two-tailed t-test, t(2) = 7.283, p = 0.0183, R^2^ = 0.9637, difference between means ± SEM = .1387 ± 0.019 with 95% CI: 0.057 0.22]). ***D1***, Violin plots with medians and quartiles, depicting kernel density of the MOC across subjects for PV particles segregated into two categories: somata + proximal dendrites, dark grey; or other dendrites not traceable to a soma, light grey. ***D2***, MOC means across species and proximity to soma classification (mean MOC marmosets soma and proximal, 0.62±0.01; marmosets other 0.47±0.02; macaques soma and proximal 0.47±0.004, macaques other 0.39±0.01; two-way ANOVA, species F_1,4_ = 293.7, p < 0.0001, explaining 55% of total variation; proximity to soma F_1,4_ = 216.9, explaining 41% of total variation, p = 0.0001; interaction F_1,4_ = 16.27, explaining 3% of total variation, p = 0.0157; Pairwise testing with Sidak’s correction confirmed significant effects within species [marm prox vs. marm other p = 0.0011, mean difference 0.1403, 95% CI: 0.089, 0.19; mac prox vs. mac other, p = 0.0098, mean difference 0.080, 95% CI: 0.029, 0.13], and between species [marm prox vs. mac prox, p = 0.0007, mean difference 0.1583, 95% CI: 0.11, 0.21; marm other vs. mac other, p = 0.0045, mean difference 0.098, 95% CI: 0.047,0.15]). ***E-F***, Confocal photomicrographs of single focal planes imaged in layer III dlPFC labeled for PV (red) and D1R (green). Images depict exemplar PV neurons that fell in the D1R strong category (D1strong, yellow arrows, **E**), or D1R negative category (D1neg, white arrowheads, **F**) for analyses in ***G-H. G***, Proportion of PV neurons falling into four D1R expression bins (D1neg, D1weak, D1moderate, D1strong) by subject. Bins were determined objectively for each individual image by binning the image’s own D1R signal intensity range. To be classified as D1neg, D1R mean grey value (MGV) in the PV neuron was less than the average D1R MGV sampled in the immunonegative “neuropil”. To be classified as D1strong, the D1R MGV of the PV neuron was above the minimum MGV of the most “strongly” labeled D1R neurons in the image, which were typically pyramidal in morphology. Intermediate D1R expression categories D1weak and D1moderate are intermediate bins between D1neg and D1strong. ***H***, Species mean for each D1R expression category (two-way ANOVA, F_3,8_ = 215.4, p < 0.0001, with Sidak’s corrected multiple comparison test, mean D1neg marmosets 25±4%, macaques 35±1%, 95% CI for mean difference: 1.33, 16.43, p = 0.022; mean D1strong marmosets 6±0.3%, macaques 3±1%, not significant). Macaques, purple; marmosets, orange. ^*^, p <0.05 using two-way ANOVA with Sidak’s corrected pairwise multiple comparisons test; n.s., not significant; D1R, dopamine 1 receptor; dlPFC, dorsolateral prefrontal cortex; Mac, macaque; Marm, marmoset; MOC, Manders’ Overlap Coefficient; PV, parvalbumin; SD, standard deviation.

Labeling was also observed in pyramidal neuron apical dendrites, confirmed by co-labeling with the cytoskeletal protein microtubule-associated protein 2 (MAP2, Figure S1). In our double-labeled PV/D1R tissue, we sampled from layer III in both species using confocal microscopy. The sampled images were analyzed in two independent ways: (1) using an automated higher throughput colocalization analysis of PV neuronal segments (Figure 3C-D), which computed the spatial extent to which PV+ processes contain colocalized PV+/D1R+ pixels; and (2) then using a manual analysis of D1R fluorescence intensity signal in PV somata (Figure 3G-H), to determine the proportion of neurons that were negative for D1R or expressed D1R weakly, moderately, or strongly.

For the colocalization analysis, we first used automatic segmentation to isolate PV neuronal processes. We then computed an index of colocalization, called the Manders’ Overlap Coefficient (MOC), to determine the proportion of each PV segment that contained colocalized D1R+/PV+ pixels. Marmoset PV segments had a higher degree of colocalization with D1R than macaque PV segments (Figure 3C). For both species, segmented particles found in the PV neuron soma and proximal dendrites had a higher degree of D1R colocalization than those in non-proximal dendrites (Figure 3D), and this disparity was greater in marmosets. The second analysis (schematized in Figure S2) binned PV neurons according to their level of D1R expression, and showed that macaques had slightly more D1R-negative PV neurons than marmosets (Figure 3E-H). This is in the same direction as the species difference in the proportions of *DRD1+ PVALB* nuclei found in our sn-RNAseq data. The trend continued with marmosets having more D1R-strong PV neurons than macaques (Figure 3H), though this was not significant.

#### Computational modeling of the effect of the species difference in D1R expression in layer III dlPFC PV neurons on distractibility during cognitive performance

### Modeling of visuospatial working memory maintenance in a microcircuit featuring structured inhibition

Given that D1R stimulation increases the intrinsic excitability of fast-spiking, putative PV neurons in macaque layer III dlPFC^**51**^, and that PV neurons shape the spatial tuning of the excitatory pyramidal population activity^**23**, **24**^, we investigated whether higher D1R expression in marmoset layer III dlPFC PV neurons may contribute to higher distractibility in cognitive tasks. We predict that similar fractions of active D1Rs on PV neurons between species—yet a higher absolute number of active D1Rs in marmosets—would cause consistently higher microcircuit inhibition in marmosets, thus suppressing task-related persistent activity within the typical, mid D1R stimulation range and increasing its susceptibility to interference by distractors. For this purpose, given the evidence for delay-related persistent activity in both marmosets^**38**^ and macaques^**14**, **15**^,we first redesigned a well-established spiking neural network model^**64**^ of visuospatial working memory maintenance in a macaque layer III dlPFC microcircuit (Figure 4A1), which allows the linking of functional microanatomy with cognitive performance. Specifically, in addition to near feature-tuned excitatory (E) Delay cells (Figure 4A2), we incorporated two new types of inhibitory Delay cells – I_*NEAR*_ and I_*OPP*_ – with near-feature and opposite-feature tuning (Figure 4A3-A4), to replace the unstructured inhibition in the original microcircuit model, because evidence suggests that putative fast-spiking, PV neurons might instead mediate such structured inhibitory motifs in primary sensory^**27-30**, **33**^ and higher order association^**25**, **26**, **31**, **32**^ cortex. Reportedly, I_*NEAR*_-mediated inhibition primarily stabilizes the amplitude of the spatially tuned population activity of excitatory Delay cells^**65-67**^, thus increasing its robustness to excitatory perturbations, like noise. In our visuospatial working memory model, such robustness could additionally translate into resistance to distractors, specifically near the target because I_*NEAR*_-mediated inhibition is strongest at and around the target position. Conversely, I_*OPP*_-mediated inhibition critically enables the competition between opposite feature-tuned Delay cell populations^**67-69**^. Accordingly, in our modeling, I_*OPP*_-mediated inhibition may confer higher resistance to far distractors by more strongly suppressing Delay E cells tuned to opposite (vs. near) positions following target presentation, compared to baseline. This increases the stimulus strength necessary to form persistent activity at the farthest positions and disrupt the target-specific persistent activity. Thus, our simulations may identify a plausible neurophysiological mechanism linking the higher D1R expression in layer III dlPFC PV neurons in marmosets with their generally recognized poorer performance in working memory maintenance tasks^**5-7**^.

**Figure 4.**
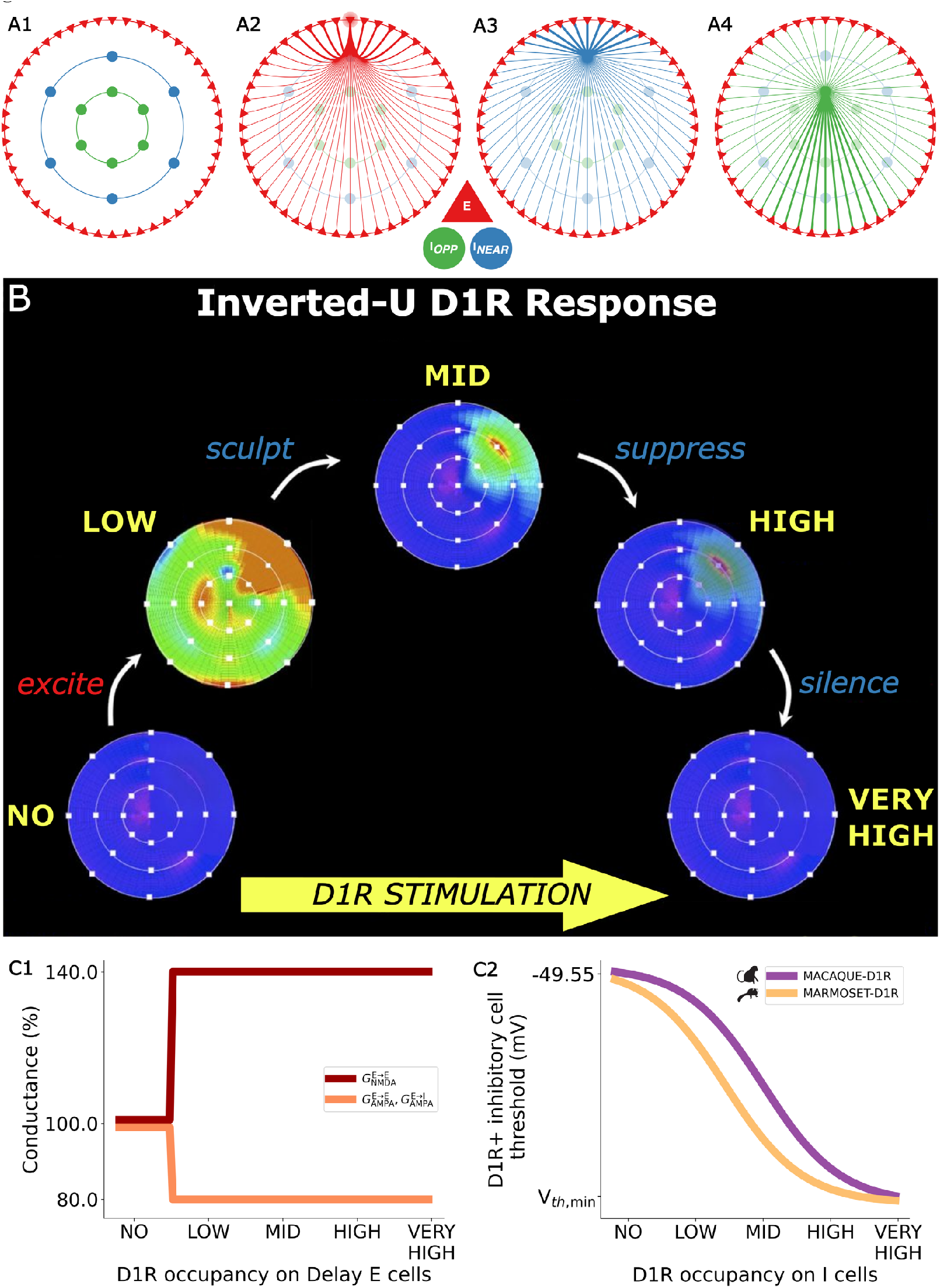
Spiking network architecture and synaptic connectivity in the visuospatial working memory model, the inverted-U relationship between D1R stimulation and persistent activity during the delay period of visuospatial working memory tasks, and microcircuit effects of D1R occupancy. ***A***, Simplified schematic representations of **(A1)** the functional architecture of the Delay cell microcircuit comprising 80% near feature-tuned E cells, 10% I_*NEAR*_ cells, and 10% I_*OPP*_ cells, with example connectivity profiles for **(A2)** an E cell to each other E cell, **(A3)** an I_*NEAR*_ cell to all E cells, and **(A4)** an I_*OPP*_ cell to all E cells, where the source cell is positioned at 0°. For each profile, a halo surrounds the cell from which the projections originate. Projection thickness indicates relative connection strength. E cells are arranged uniformly on a ring, in functional feature space, according to the angle of the visual stimuli for which they fire the most. Inhibitory cells are also arranged uniformly on a ring, based on the angular position of the E cells from which they receive the strongest excitation. The microcircuit features E→E, E→I, I→E, and I→I synapses. E cells send their strongest glutamatergic projections to all cells at the same angle as themselves. I_*NEAR*_ cells send their strongest GABAergic projections to all cells also at the same angle as themselves, while I_*OPP*_ cells – to all cells positioned 180° away from them on the ring. ***B***, Diagram illustrating the idealized inverted-U response, characterized by the delay-period activity during visuospatial working memory maintenance in macaques under varying D1R stimulation conditions (adapted with permission from Arnsten *et al*.^**42**^). At one extreme, no D1R stimulation means E cells cannot support persistent activity during the delay period, with only transient activity recorded during stimulus application. In the increasing phase of the inverted-U, low (suboptimal) D1R stimulation induces erroneous (here specifically, noise-evoked and/or spatially diffuse) persistent activity due to disinhibition. At the peak of the inverted-U, mid (optimal) D1R stimulation sculpts spatially precise, noise- and distractor-resistant persistent activity. Crucially, in the decreasing phase of the inverted-U, high (supraoptimal) D1R stimulation suppresses E cells, decreasing their firing rates, and thus increasing the susceptibility of the persistent activity for target stimuli to disruptions by distractors. Finally, at the other extreme of the inverted-U, very high D1R stimulation silences E cells completely, precluding the formation of persistent activity, again with only transient activity being recorded during stimulus application. ***C1***, Step-like relationships between D1R occupancy and (1) *G*_E→E, NMDA_ and (2) *G*_E→E, AMPA_ and *G*_E→I, AMPA_, as percentages of their respective baseline values. From no to low D1R occupancy, specifically at *x* = 0.125, we increased *G*_E→E, NMDA_ by 40% and decreased *G*_E→E, AMPA_ and *G*_E→I, AMPA_ by 20%. We applied the same D1R modulations of *G*_E→E, NMDA_, *G*_E→E, AMPA_, and *G*_E→I, AMPA_, in the marmoset-D1R model and the macaque-D1R model. ***C2***, Sigmoidal relationship between D1R occupancy and the species-D1R model-specific firing thresholds of the D1R+ inhibitory cells across the microcircuit models (and the different networks of the working memory maintenance model). We decreased the firing thresholds of D1R+ inhibitory cells from their baseline (V_*th*, base_) of -49.55 mV at no D1R occupancy, through the low, mid, and high levels, and to a minimum (V_*th*, min_) at the very high level. V_*th*, min_ differed across networks, as listed in Table 4 in the Methods.

### D1R expression in Delay E and I_NEAR_ cells facilitates the inverted-U response

We considered three cell type-specific D1R expression patterns, and explored which could reproduce the inverted-U relationship between D1R stimulation and persistent firing activity observed experimentally during the delay period of visuospatial working memory tasks^**42**, **43**, **45**^ (Figure 4B). Specifically, we built three networks in which Delay E cells always expressed D1Rs, while D1R expression in the inhibitory Delay cells differed. D1Rs were expressed in Network 1 by I_*NEAR*_ cells; in Network 2 by I_*OPP*_ cells; and in Network 3, by both I_*NEAR*_ and I_*OPP*_ cells. We considered D1R stimulation in terms of D1R occupancy, the fraction of DA-bound D1Rs. We defined five main D1R occupancy levels – no (0), low (0.25), mid (0.50), high (0.75) and very high (1) – on a normalized scale ranging from 0 to 1, and evaluated network performance in a visuospatial working memory maintenance task. During task trials, we briefly applied a target stimulus at 0° and, later, a distractor stimulus at 180°, the most representationally dissimilar position. We varied the strength of stimuli to identify what strength evoked the most reliable inverted-U response. Too weak stimuli will never elicit persistent activity, while too strong stimuli will always cause distractibility. To reproduce the distinct persistent activity regimes of the inverted-U in a macaque-D1R model of covert working memory maintenance, we implemented a set of D1R modulations of synaptic and cellular parameters. Briefly, from no to low D1R occupancy, we increased the NMDAR conductance at E→E synapses^**70**, **71**^ (*G*_E→E, NMDA_) to facilitate a transition from a transient to a persistent activity regime and (2) decreased AMPAR conductances at E→E and E→I synapses^**70**, **72**^ (*G*_E→E, AMPA_, *G*_E→I, AMPA_) to reduce the impact of noise-evoked activations of E cells on glutamatergic transmission to other E and I cells (Figure 4C1). We also decreased the firing thresholds of D1R+ inhibitory cells to increase their intrinsic excitability and respectively sculpt, suppress, and silence persistent activity at mid, high, and very high D1R occupancy (Figure 4C2).

Network 1 (D1R+ E, I_*NEAR*_) reproduces the inverted-U over a range of stimulus strengths between ∼0.15-0.30 pA with high reliability (max. fraction of complete inverted-Us of 99.1% at 0.20 pA) (Figure S4). Specifically, no and very high D1R occupancies result in transient activations in response to the stimuli due to, respectively, insufficient recurrent excitation and overinhibition (Figure 5A1 and 5E1). Low D1R occupancy elicits a disinhibited, erroneous regime, prone to spatially diffuse and noise-evoked persistent activity at non-stimulus positions (Figure 5B1). Mid D1R facilitates a spatially precise, noise- and distractor-resistant persistent activity regime, suggesting the network represents the memory more accurately during the delay period due to optimal inhibition (Figure 5C1). Notably, high D1R occupancy induces a distractible regime where persistent activity forms around the target position during the delay period but then jumps to the position of the distractor when it appears (Figure 5D1). In contrast, Network 2 (D1R+ E, I_*OPP*_) cannot reproduce the inverted-U (Figure S4), particularly the distractible persistent activity expected at high D1R occupancy. Network 3 (D1R+ E, I_*NEAR*_, I_*OPP*_) reproduces the inverted-U, like Network 1, but over a narrower range between ∼0.20-0.30 pA, and less reliably (max. fraction of complete inverted-Us of 67.7% at 0.25 pA) (Figure S4). Overall, microcircuits featuring D1R+ E and I_*NEAR*_ cells (Network 1 or 3) appear sufficient to reproduce the inverted-U relationship between D1R stimulation and *in vivo* delay-period neuronal firing activity in visuospatial working memory tasks, unlike a microcircuit featuring only D1R+ E and I_*OPP*_ cells (Network 2). Our data suggest that D1R+ I_*NEAR*_ cells may represent D1R+ PV neurons because specifically D1R+ I_*NEAR*_ cells contributed, alongside D1R+ E cells, to the inverted-U response by increasing their intrinsic excitability with D1R stimulation, which may occur only for D1R+ fast-spiking, putative PV neurons^**51**^. This corroborates indications^**73-75**^ that PV neurons provide near-feature inhibition in microcircuits.

**Figure 5.**
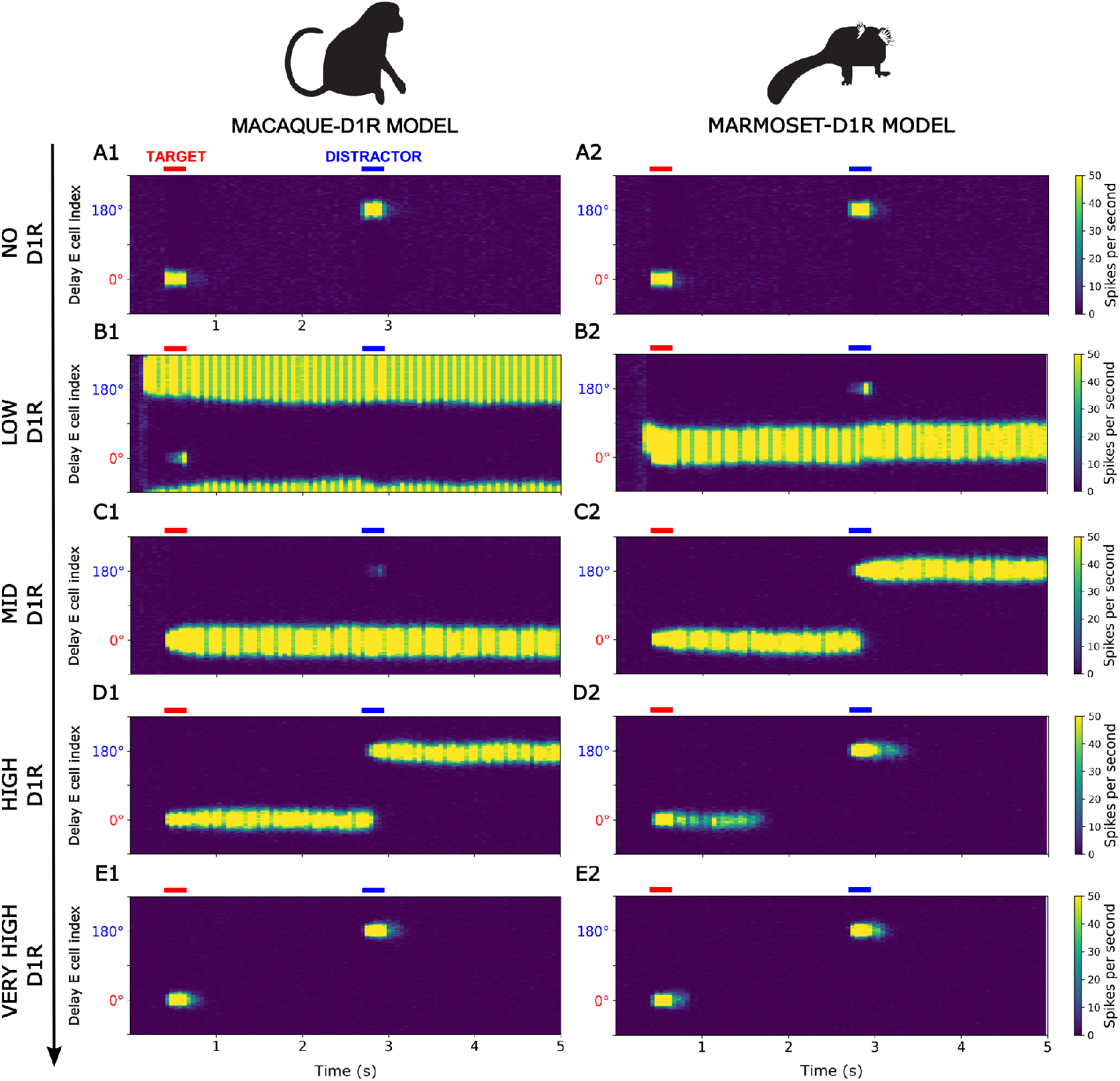
Distractible, decreasing phase of inverted-U shifted to typical, mid D1R occupancy in the marmoset-D1R model. Rastergrams represent the color-coded, unmanipulated spike time series of Delay E cells in Network 1 for the macaque-D1R model **(A1-E1)** and the marmoset-D1R model **(A2-E2)**, during example five-second trials of the simulated visuospatial working memory maintenance task across the five main D1R occupancy levels **(A-E)**. Delay E cells are indexed by the angular position from 0° to 360° of their preferred stimuli. The strength of the stimuli is 0.20 pA. For visualization purposes, the y-axis wraps around 270° to accentuate the target (0°) and distractor (180°) positions, and the color scale is capped to reduce the dominance of outliers in the heatmap, such that any firing rate of >50 Hz per second per neuron is plotted in the same color as firing rates of 50 Hz.

### Most plausible visuospatial working memory performance in microcircuit with D1R+ Delay E and I_NEAR_ cells, and D1R-Delay I_OPP_ cells

To further evaluate the effect of inhibitory cell-type-specific D1R expression on visuospatial working memory maintenance in Networks 1-3 (Figure S5), we simulated the effects of distractors applied at different positions. We focused on Networks 1 and 3, specifically at the typical, mid D1R occupancy level, because Network 2 cannot reproduce the inverted-U for distractors even at 180°. Whereas Network 3 performed poorly due to overwhelming distractibility or errors across distractor positions, with distractor resistance predominating only around 180°, Network 1 was overwhelmingly distractor-resistant at intermediate-to-far positions (112.5°-180°) but also significantly at near positions (45°). We also examined two alternative networks featuring Delay E cells and only, respectively, Delay I_*NEAR*_ or I_*OPP*_ cells (all D1R+) (Figure S6). The I_*NEAR*_-only network exhibited significant resistance to near distractors (45°) but also overwhelming distractibility and errors across distractor positions. Conversely, the I_*OPP*_-only network could overwhelmingly resist far distractors (112.5°-180°), while being highly distractible for near distractors (45°-90°). Furthermore, expectedly, the I_*OPP*_-only network could not be distracted by far stimuli at high D1R occupancy (135°-180°). Overall, I_*NEAR*_ and I_*OPP*_ cells enable near and far distractor resistance, respectively, and both are necessary for effective microcircuit operations, with D1R expression in only E and I_*NEAR*_ cells (Network 1) enabling the most plausible performance. These results match experimental observations in a similar delayed-response visuospatial working memory task^**76**^, which suggested that macaque dlPFC microcircuitry provides some resistance to near distractors (45°) and strong resistance to far distractors (135°, 180°).

### Distractibility induced in the typical, mid D1R occupancy range in a marmoset-D1R model of covert working memory maintenance

To examine how the higher D1R expression in the layer III dlPFC PV neurons of marmosets could affect their distractibility in tasks requiring working memory maintenance, we integrated our neuroanatomical results from marmosets into the most functionally plausible network, namely Network 1, of the working memory maintenance model. In this manner, we established a marmoset-D1R model of the working memory maintenance microcircuit, which differed from the macaque-D1R model only in the higher D1R expression in the putative PV cells and, more importantly, in its neurophysiological consequences. Specifically, given the cross-species difference in D1R expression in PV neurons (Figure 3), we hypothesize that more D1Rs are DA-bound on PV neurons in marmosets compared to macaques, assuming similar extracellular DA concentration in dlPFC across conditions between the two species. Essentially, the same D1R occupancy signifies greater D1R stimulation of marmoset dlPFC PV neurons. Thus, since D1R+ I_*NEAR*_ cells likely represent D1R+ PV neurons, we shifted the midpoint of the I_*NEAR*_ threshold sigmoid from the reference value of 0.50 in the macaque-D1R model, on the normalized D1R occupancy scale, to 0.36 in a marmoset-D1R model of covert working memory maintenance, based on the difference (0.14) in mean MOCs between the species (Figure 4C2).

As in the macaque-D1R model, the marmoset-D1R model shows only transient activity for no and very high D1R occupancy (Figure 5A2 and 5E2). For low D1R occupancy, the marmoset-D1R model overwhelmingly exhibits erroneous spatially diffuse, noise-evoked persistent activity (Figure 5B2), but with a lower incidence of 85.4% compared to the macaque-D1R model’s 97.3% (Figure 6A-B). Crucially, for mid D1R occupancy (Figure 5C2), the marmoset-D1R model demonstrates distractible persistent activity, which can be verified experimentally, with an incidence of 87.2%, unlike the macaque-D1R model’s 99.7% distractor-resistant persistent activity (Figure 6A-B). Finally, at high D1R occupancy, the marmoset-D1R model already transitions to the transient activity regime (Figure 5D2), which occurs at a higher D1R occupancy in the macaque-D1R model (Figure 6A-B). Overall, the modeling predicts that the higher D1R expression in PV neurons (putative I_*NEAR*_ cells) in marmoset layer III dlPFC shifts the apex and decreasing phase of the inverted-U to lower D1R occupancy levels in marmosets, compared to macaques, since the optimal PV-mediated inhibition is closer to the peak NMDAR-mediated excitation. Thus, the typical, mid D1R occupancy range supports spatially precise, noise- and distractor-resistant persistent activity in macaques, while inducing distractible persistent activity in marmosets.

**Figure 6.**
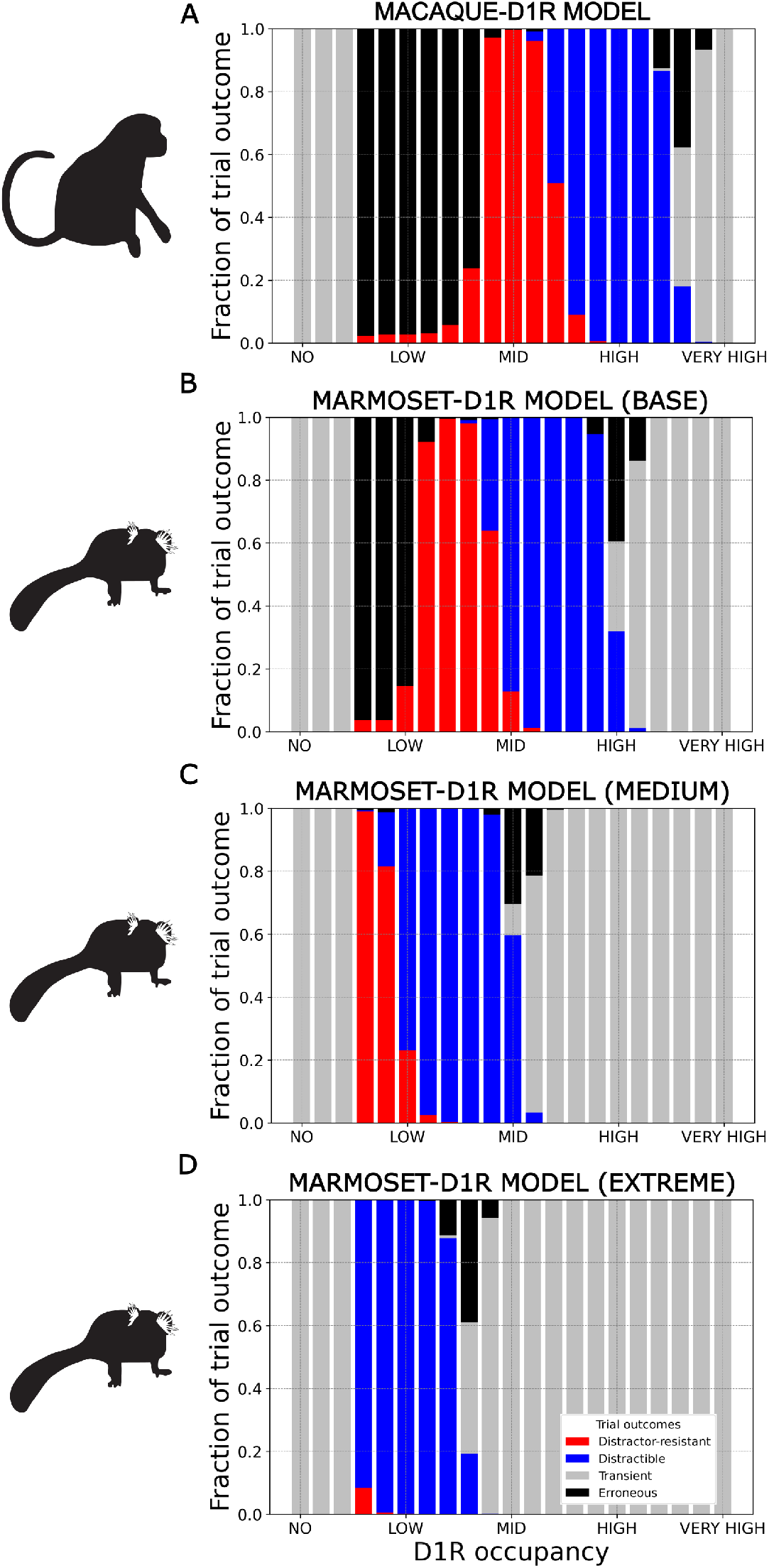
Progressively leftward shift of inverted-U with greater D1R stimulation. Fractions of classified outcomes of 1000 trials across the granulated D1R occupancy scale in the visuospatial working memory maintenance task with distractors presented only at 180° to Network 1 for the macaque-D1R model and the three cases of V_*th*_ sigmoid midpoint shift for the marmoset-D1R model. The strength of the stimuli is 0.20 pA. In the no D1R occupancy range, both species-D1R models **(A-D)** show transient activity. In the low D1R occupancy range, the macaque**-**D1R model **(A)** exhibits predominantly erroneous persistent activity, while the base marmoset**-**D1R model case **(B)** transitions from the erroneous to the noise- and distractor-resistant persistent activity regime, the medium marmoset-D1R model case **(C)** transitions from the noise- and distractor-resistant to the distractible persistent activity regime, and the extreme marmoset-D1R model case **(D)** exhibits predominantly distractible persistent activity. In the mid D1R occupancy range, the macaque-D1R model **(A)** overwhelmingly exhibits noise- and distractor-resistant persistent activity. The base marmoset-D1R model case **(B)** transitions to the distractible persistent activity regime, with a decreasing fraction of noise- and distractor-resistant persistent activity. The medium marmoset-D1R model case **(C)**, transitions to transient activity, with a decreasing fraction of distractible persistent activity and a significant fraction of erroneous persistent activity (indicating persistent activity that is disrupted before the readout due to overinhibition). The extreme marmoset-D1R model case **(D)**, which predominantly exhibits transient activity. In the high D1R occupancy range, the macaque-D1R model **(A)** reliably exhibits a distractible persistent activity regime, while the base marmoset-D1R model case **(B)** transitions from the distractible persistent activity to the transient activity regime, with a significant fraction of erroneous persistent activity, and the medium and extreme marmoset**-**D1R model cases **(C-D)** exhibit transient activity. Finally, in the very high D1R occupancy range, both species-D1R models **(A-D)** settle into a transient activity regime. Notably, as the shift increases across the marmoset-D1R model cases, the range of D1R occupancies which can support persistent activity decreases.

However, because differences in the MOC-measured D1R expression in I_*NEAR*_ cells may not correspond directly to differences in their D1R stimulation and the resulting effects on microcircuit function (e.g., due to limitations of the D1R measurement method or differences in D1R binding, coupling efficiency, etc.), we examined larger sigmoid midpoint shifts (Figure 6C-D). Notably, a shift of 0.375 was approximately the largest that still supports (distractible) persistent activity at the typical, mid D1R occupancy in the marmoset-D1R model before the working memory maintenance microcircuit enters a transient activity regime. Thus, shifts between ∼0.14-0.375 would capture the cross-species differences in distractibility, while allowing for plausible microcircuit functioning in the typical, mid D1R stimulation range.

### Computational modeling of the effect of the species difference in D1R expression in layer III dlPFC PV neurons on distractibility during visual fixation

Finally, we examined how the higher D1R expression in layer III dlPFC PV neurons of marmosets could affect their distractibility in visual fixation tasks requiring overt sustained attention. Notably, although overt sustained attention may share neurophysiological mechanisms underlying persistent activity with covert working memory maintenance^**14-21**^, evidence suggests that the specific neuronal substrates that support appropriately tuned, noise- and distractor-resistant persistent activity in the visuospatial working memory microcircuit may only partially overlap with those recruited in our visual fixation task^**11-13**^ (Figure 1B). In fact, the shared neuronal substrates between the two functions may only support the selective allocation of attention to specific items within the working memory space. Furthermore, our model of covert working memory maintenance did not simulate visual fixation (a component of the real-world oculomotor delayed-response task). For these reasons, we further designed a novel spiking network model of overt sustained attention in a primate dlPFC microcircuit (Figure 7A1). Specifically, we implemented Cue cells, including near feature-tuned E and I_*NEAR*_ cells arranged on a ring, which exhibit only transient activations for the distractor stimuli in our visual fixation task. Importantly, we also implemented Fixation cells, selective for the center of the visual feature space, which included identically tuned Fixation Rule E cells (Figure 7A2), Fixation I_*NEAR*_ cells, and Fixation I_*OPP*_ cells. Fixation Rule E cells maintain the rule to fixate overtly on the central point and may represent layer III pyramidal neurons because the former can sustain persistent activity^**17**^ (supported by layer III microcircuitry), after the removal of the central point that drove activity externally. Critically, Fixation I_*NEAR*_ cells mediate local feedforward inhibition, driven by the excitatory Cue cells, onto the Fixation Rule E cells (Figure 7A3), in addition to stabilizing the firing rates of Fixation Rule E cells through feedback inhibition. Driven by the Fixation Rule E cells (Figure S7), the Fixation I_*OPP*_ cells mediate local top-down inhibition onto the Cue E cells (Figure 7A4), which suppresses peripheral visual distractors—a central aspect of sustained attention^**77**^.

**Figure 7.**
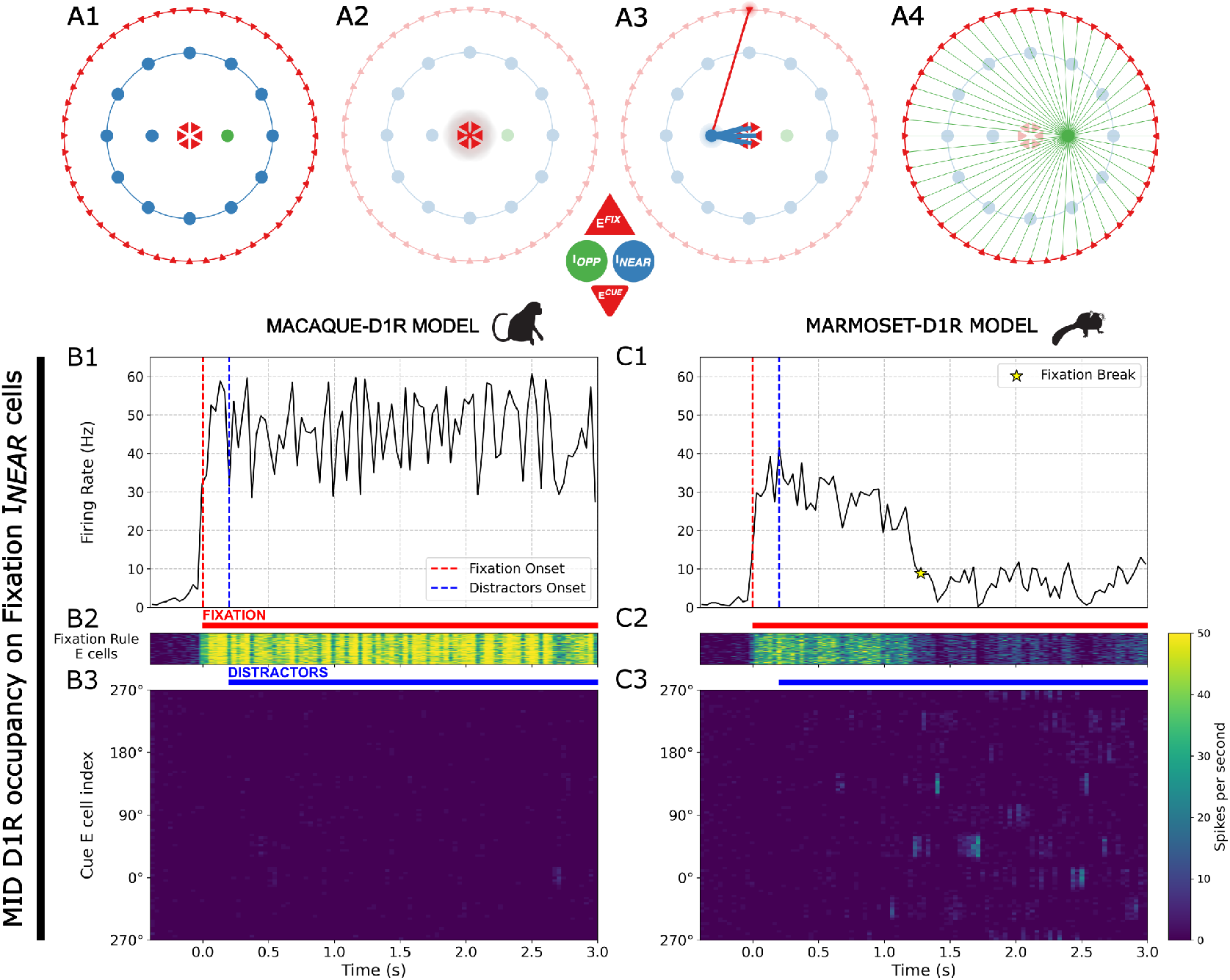
Spiking network architecture and synaptic connectivity in the visual fixation model, and example trials in the simulated task. Simplified schematic representations of **(A1)** the functional architecture of the microcircuit, comprising 71.11% near feature-tuned Cue E (E^*CUE*^) cells, 17.78% Cue I_*NEAR*_ cells, 8.89% identically tuned Fixation Rule E (E^*FIX*^) cells, 1.11% Fixation I_*NEAR*_ cells, and 1.11% Fixation I_*OPP*_ cells, **(A2)** all Fixation Rule E cells fully connected with each other with equal synaptic strengths, **(A3)** the local feedforward inhibitory motif where an example Cue E cell positioned at 0° excites a Fixation I_*NEAR*_ cell, which in turn inhibits the Fixation Rule E cells, **(A4)** the local top-down inhibitory motif where an example Fixation I_*OPP*_ cell, driven by the Fixation Rule E cells (Figure S7), inhibits the Cue E cells. Projection thickness indicates relative connection strength. Cue E cells are arranged uniformly on a ring, in functional feature space, according to the angle of the visual distractor stimuli for which they fire the most. Cue I_*NEAR*_ cells are also arranged uniformly on a ring, based on the angular position of the Cue E cells from which they receive the strongest excitation. Fixation Rule E cells represent the central fixation rule and have no angular position selectivity. The Fixation I_*NEAR*_ cells are co-tuned with the Fixation Rule E cells. Histograms represent the average firing rate in the Fixation Rule E population in the macaque-D1R model **(B1)** and the marmoset-D1R model **(C1)**, during example three-second trials of the simulated visual fixation task at the typical, mid level of D1R occupancy on Fixation I_*NEAR*_ cells. The red dashed line at 0 seconds indicates the onset of the fixation cue. The blue dashed line at 0.2 seconds indicates the onset of the distractor cues. Rastergrams represent the color-coded, unmanipulated spike time series of Fixation Rule E cells and Cue E cells in the macaque-D1R model **(B2-3)** and the marmoset-D1R model **(C2-3)**. Cue E cells are indexed by the angular position from 0° to 360° of their preferred stimuli. The strength of the stimuli is 0.20 pA. For visualization purposes, the y-axis wraps around 270° to be consistent with Figure 5, and the color scale is again capped to reduce the dominance of outliers in the heatmap, such that any firing rate of >50 Hz per second per neuron is plotted in the same color as firing rates of 50 Hz.

Having established in our model of covert working memory maintenance that D1R+ I_*NEAR*_ cells likely represent D1R+ layer III dlPFC PV neurons, for our purposes here, we considered only one cell type-specific D1R expression pattern where only Fixation I_*NEAR*_ cells expressed D1Rs. First, we explored how an inverted-U relationship between D1R stimulation and persistent activity would manifest during visual fixation. We again considered D1R stimulation in terms of D1R occupancy, the fraction of DA-bound D1Rs, but only on Fixation I_*NEAR*_ cells. We produced distinct persistent activity regimes of the Fixation Rule E cells specifically through D1R modulation of the firing thresholds of Fixation I_*NEAR*_ cells to increase their intrinsic excitability and respectively sculpt, suppress, and silence persistent activity at mid, high, and very high D1R occupancy (Figure 4C2). We decreased the firing thresholds of Fixation I_*NEAR*_ cells in the macaque-D1R and marmoset-D1R models of overt sustained attention across the scale of D1R occupancy (on Fixation I_*NEAR*_ cells), as described in Figure 4C2. In particular, we implemented the I_*NEAR*_ threshold parameters from Network 1 of the working memory maintenance model to ensure the microcircuit models are comparable within the same D1R modulation framework. Finally, we evaluated model performance in a visual fixation task requiring sustained attention, like our behavioral task performed by marmosets and macaques. Briefly, we initiated task trials with a fixation stimulus applied continuously to the Fixation Rule E cells until trial offset. Soon after presenting the fixation stimulus, we applied a stream of short peripheral distractor stimuli to the Cue E cells, randomly sampled from nine angular positions on the ring. Thus, overall, our simulations may explain our measures of behavioral performance in the visual fixation task using the neurophysiological mechanism of distractibility identified in our model of covert working memory maintenance.

### Higher distractibility and more variable fixation duration in the typical, mid D1R occupancy range in marmoset-D1R model of overt sustained attention

We focused on the typical, mid D1R occupancy level on Fixation I_*NEAR*_ cells, where the differences in fixation performance should be most apparent. The marmoset-D1R model demonstrates distractible persistent activity (Figure 7B), in 99.8% of trials (Figure 8A1). This contrasts with the macaque-D1R model’s distractor-resistant persistent activity (Figure 7C), in 99.7% of trials (Figure 8A2). We performed a permutation test to assess differences in mean fixation durations between the marmoset-D1R and macaque-D1R models within the mid D1R occupancy range. Under the null hypothesis of no difference, 10,000 permutations were conducted by randomly shuffling species-D1R model labels across trials. The observed difference in means of 1.31 seconds (*M*_MARM-D1R_ = 1.31 seconds, *SD*_MARM-D1R_ = 1.04 seconds; *M*_MAC-D1R_ = 2.62 seconds, *SD*_MAC-D1R_ = 0.75 seconds) was highly significant (p < 0.001), indicating that mean fixation duration in the marmoset-D1R model was significantly shorter than in the macaque-D1R model (Figure 8B2). Thus, the visual fixation model closely reproduced the cross-species differences in distractibility and fixation duration in our behavioral results. This suggests that the higher D1R expression in layer III dlPFC PV neurons in marmosets could also contribute to their higher distractibility and shorter fixation duration during overt sustained attention via more prevalent disruptions of persistent activity at the typical, mid range of D1R occupancy, like in our working memory maintenance model. Additionally, fixation durations in the marmoset-D1R model were more variable than in the macaque-D1R model (coefficients of variation (CVs): 0.80 (79.5%) for the marmoset-D1R model and 0.29 (28.8%) for the macaque-D1R model). To assess whether this difference in variability was statistically significant, we conducted a permutation test on the CVs with 10,000 permutations. The observed difference in CVs (ΔCV = 0.51) was highly significant (p < 0.0001). Thus, our modeling further suggests that higher D1R expression in layer III dlPFC PV neurons may contribute to more variable cognitive performance.

**Figure 8.**
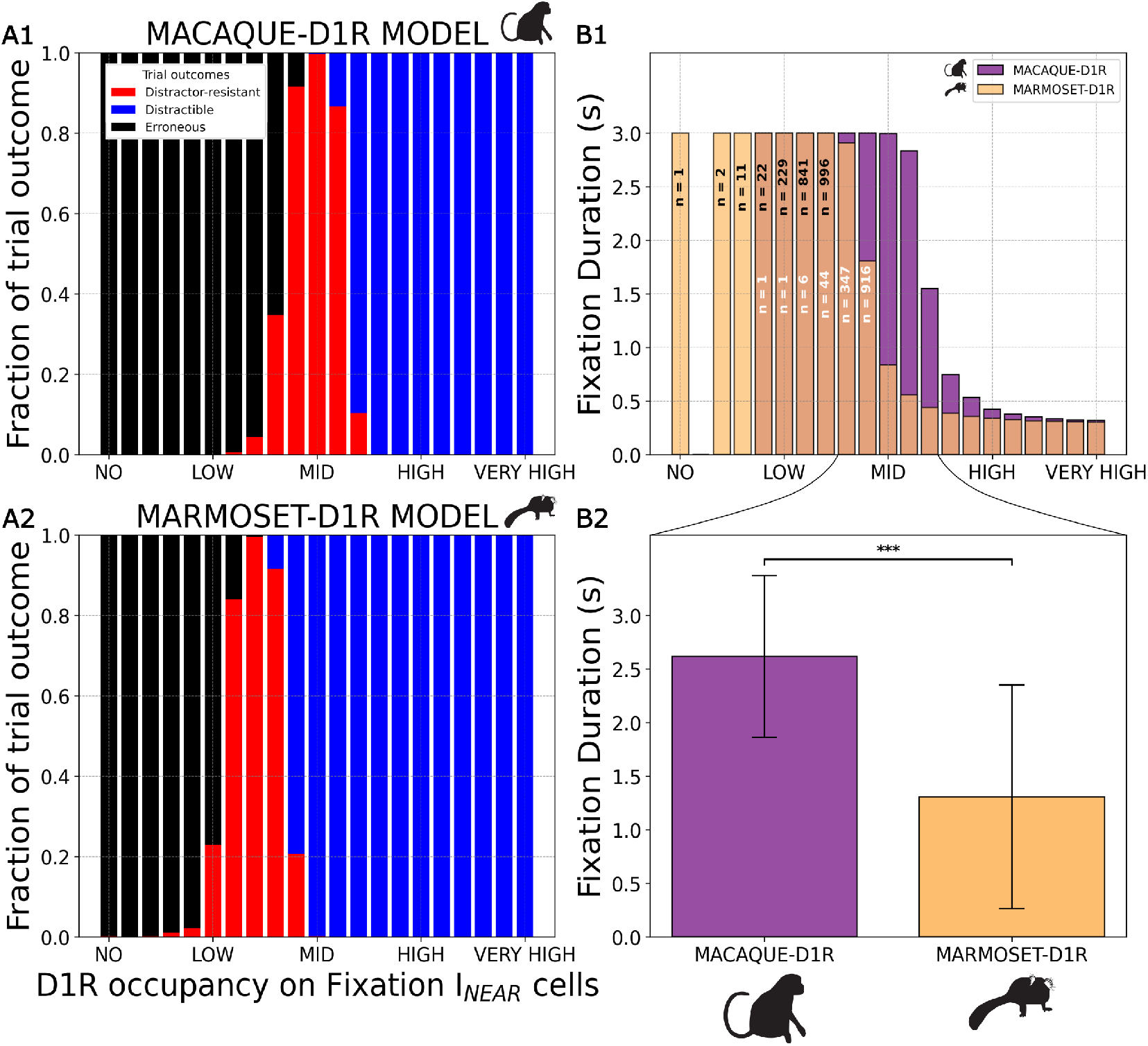
Marmoset-D1R and macaque-D1R models of sustained attention reproduce differences in behavioral performance during visual fixation between the species. ***A***, Fractions of classified outcomes of 1000 trials across the granulated scale of D1R occupancy on Fixation I_*NEAR*_ cells for each species-D1R model. The strength of the stimuli is 0.20 pA. In the no and low ranges of D1R occupancy on Fixation I_*NEAR*_ cells, the macaque**-**D1R model **(A1)** exhibits predominantly erroneous persistent activity, while the marmoset**-**D1R model **(A2)** transitions from the erroneous to the noise- and distractor-resistant persistent activity regime. In the mid D1R occupancy range, the macaque-D1R model overwhelmingly exhibits noise- and distractor-resistant persistent activity, while the marmoset-D1R model transitions to the distractible persistent activity regime, with a decreasing fraction of noise- and distractor-resistant persistent activity. Finally, in the high and very high D1R occupancy ranges, both species-D1R models reliably exhibit a distractible persistent activity regime. ***B***, Mean fixation duration in the 1000 trials across the granulated scale of D1R occupancy on Fixation I_*NEAR*_ cells, and within the mid D1R occupancy range, in the visual fixation task for each species-D1R model. Until the peak of the optimal level of D1R occupancy on Fixation I_*NEAR*_ cells, the mean fixation duration from fixation cue onset was 3 seconds, indicating complete trials without fixation breaks, for both species-D1R models. Afterwards, the mean fixation duration began to decrease to about 0.30 seconds at the very high D1R occupancy level for both species-D1R models **(B1)**. The mean fixation duration for the marmoset-D1R model began decreasing at a lower D1R occupancy relative to that for the macaque-D1R model. The mean fixation durations for the marmoset-D1R model are overlaid onto those for the macaque-D1R model with transparent coloring. Notably, when estimating the mean fixation durations, we excluded trials where the species-D1R models exhibited erroneous persistent activity because these involved noise-evoked activations due to disinhibition in the Fixation Rule E cell population before the onset of any external stimuli (fixation or distractor cues). The white and black labels in the bar graphs in **B1** indicate the number of trials out of 1000 from which the mean fixation duration was estimated for a given D1R occupancy in respectively the macaque-D1R and the marmoset-D1R models. For bars without labels, all 1000 trials for the given D1R occupancy level were non-erroneous. Finally, we computed and compared the mean fixation durations between the species-D1R models within the mid D1R occupancy range (**B2**) by pooling the non-erroneous trials for each species-D1R model and performing a permutation test. Error bars depict ± SD. ^***^, p < 0.001.

## Discussion

We showed that marmosets are more distractible than macaques in a visual fixation task. Furthermore, we found that marmosets have higher *DRD1* expression in *PVALB* neurons than macaques, with marmoset levels comparable to those in mice. In contrast, humans showed the lowest levels. We then confirmed *in situ* that layer III dlPFC PV neurons exhibit higher D1R expression in marmosets compared to in macaques. Finally, our microcircuit models of covert working memory maintenance and overt sustained attention suggested that higher D1R expression in layer III dlPFC PV neurons may shift the distractible persistent activity regime of the inverted-U response leftward. In particular, in the macaque-D1R models, the typical, mid range of D1R occupancy optimized Delay and Fixation Rule E cell firing. In contrast, in the marmoset-D1R models, the same D1R occupancy range suppressed and disrupted task-related persistent activity, causing more distractibility, like we observed in our marmosets. To our knowledge, this is the first study to examine how D1R expression differences in PV neurons may affect cognitively engaged, non-human primate dlPFC microcircuits.

Longstanding electrophysiological and behavioral evidence in marmosets and macaques has implicated DA in distractor resistance during working memory maintenance and sustained attention^**39**, **40**, **78-80**^, but specific microcircuit mechanisms remain an active research area. D1R prevalence on the dendritic spines of dlPFC pyramidal neurons positions them to enhance NMDAR actions in the synapse and, thus, strengthen recurrent excitation to facilitate distractor-resistant persistent activity in Delay cell microcircuits^**42**, **76**, **81**^. D1Rs may further ensure appropriate tuning of persistent activity by attenuating noisy or task-irrelevant microcircuit inputs via the opening of K+ channels on spines^**42**, **46**, **82**^, which affects distractor resistance. Another mechanism by which D1R stimulation may affect distractor resistance is through its modulation of cortical inhibition. Several microcircuit models of association cortex have already incorporated I_*NEAR*_-like^**83-86**^, and also I_*OPP*_-like^**32**, **65**, **67**, **68**^, inhibition to account for the growing evidence for these structured inhibitory motifs^**24-26**, **30-32**, **87**^, thus succeeding early attempts that featured only unstructured^**64**, **88-92**^ inhibition. However, despite the long history of simulating D1R effects on dlPFC-mediated cognition^**71**, **93-99**^, to our knowledge, no study has investigated how D1R modulation of structured inhibition affects higher cognitive functions. In rodents and macaques, D1Rs have been shown to increase inhibition through fast-spiking^**49-51**^, non-fast spiking^**100**^, or vasoactive intestinal peptide-expressing (VIP) inhibitory neurons^**101**, **102**^ Here, we simulated D1R modulation of inhibition strictly based on evidence in macaque layer III dlPFC that D1Rs excite fast-spiking, putative PV neurons^**51**^.

Specifically, given scant evidence about the organization and function of macaque dlPFC microcircuitry supporting sustained attention, or of marmoset dlPFC microcircuitry in general, we first differentiated I_*NEAR*_ and I_*OPP*_ cells in a classic working memory maintenance model based on knowledge about the dlPFC microcircuit organization and function that supports working memory maintenance in macaques. We next explored how three inhibitory cell-type-specific D1R expression profiles impact working memory maintenance. We found that a microcircuit that features excitatory and both inhibitory Delay cell types and where D1Rs modulate E and I_*NEAR*_ cells (putative D1R+ PV neurons) can reproduce the inverted-U relationship between D1R stimulation and persistent activity observed in macaques *in vivo*^**45**^, and also capture the distractor distance-dependent pattern of visuospatial working memory performance measured behaviorally in macaques^**76**^. Hence, optimal D1R stimulation may principally facilitate stabilization of the amplitude of the spatially tuned population activity of excitatory Delay cells^**22**, **66**, **67**^, and thus also possibly promote near distractor resistance.

On the other hand, the competition between opposite feature-tuned excitatory Delay cell populations, and thus far distractor resistance, may be less dependent or independent of D1R modulation because microcircuits featuring D1R+ I_*OPP*_ cells, which increase their excitability with D1R stimulation, seem functionally implausible. In particular, Network 2 (and the I_*OPP*_-only network) in our working memory maintenance model could not reproduce the distractible regime of the inverted-U for distractors at 180°, likely because when D1R stimulation increases the strength of I_*OPP*_-mediated inhibition enough to disrupt target-specific persistent activity, at far distractor positions the I_*OPP*_–mediated inhibition becomes even stronger, precluding distractor-specific persistent activity from forming. Also, Network 3 exhibited overwhelming distractibility across distractor positions at mid D1R occupancy in the macaque-D1R model, possibly due to overinhibition caused by twice as many I cells increasing their excitability, compared to Network 1. However, due to the combinatorial explosion of possibilities, we did not explore other more intricate inhibitory cell-type-specific D1R expression patterns, microcircuit configurations, etc., which could allow D1Rs to modulate competition between excitatory Delay cell populations while preserving plausible microcircuit function. Thus, while the role of D1Rs in the stabilization of persistent activity is reasonably supported by our modeling findings, evidence about D1R involvement in competitive dynamics is inconclusive.

By integrating our anatomical results into our most functionally plausible microcircuit configuration in the working memory maintenance model (Network 1), we propose that differential D1R modulation of layer III dlPFC PV neurons may underlie some cognitive differences between marmosets and macaques. Specifically, the higher D1R expression in layer III dlPFC PV neurons in marmosets may contribute to their higher distractibility by shifting the distractible persistent activity regime of the inverted-U response to the typical, mid D1R occupancy range. Conversely, the same level of DA stimulation in macaques supports appropriately tuned and distractor-resistant persistent activity for working memory maintenance. Interestingly, the D1R occupancy range that induces erroneous persistent activity in the increasing phase of the inverted-U contracted in the marmoset-D1R model, predicting that marmosets are less susceptible to internal distractions (context-irrelevant mental representations, formed without sensory input, that overtake network dynamics). Overall, by only modeling a higher D1R expression in putative PV neurons in what is essentially a layer III dlPFC microcircuit largely based on macaque research, we reproduced the higher behavioral distractibility of marmosets compared to macaques in working memory maintenance tasks^**5-7**^.

Importantly, a recent model suggested that D1R modulation of somatostatin (SST; similar to calbindin (CB) in primates) and VIP (similar to calretinin (CR) in primates) inhibitory neurons also affects cognitive performance^**99**^. The I_*OPP*_ cells in our study may represent D1R-/PV+ neurons, perhaps about a third of the total layer III dlPFC PV population according to our manual anatomical analysis (Figure 3H). Alternately, SST/CB neurons have been implicated in opposite feature-tuned inhibition^**32**, **69**, **99**, **103**^ and can display fast spikes^**104-106**^, like PV neurons. Since CB neurons target the distal dendritic trees of pyramidal neurons^**107**^, theoretical studies^**99**, **103**^ have suggested they could dampen inputs into the distal dendrites and thus attenuate the effect of noise or distracting stimuli on network dynamics. However, the identities of distal dendrite inputs are unclear and likely not all task-irrelevant. Moreover, both I_*NEAR*_ and I_*OPP*_ cells in our modeling synapse on the E cell soma because our E cells lack dendritic compartments, questioning a clear correspondence between I_*OPP*_ cells and SST/CB neurons, and supporting a correspondence with PV neurons, which target the perisomatic region^**107**^. Finally, our modeling lacks disinhibitory VIP/CR neurons, which perform permissive gating for pertinent incoming signals^**103**, **107-109**^. The CR neuron population has expanded in primates^**110**^, and CR neurons are more prominent in the dlPFC compared to earlier parts of the dorsal stream^**104**^, suggesting their prominent role in higher cognition. Future studies should explore species differences in D1R modulation of CB and CR neurons, which are also critical components of cortical microcircuits.

We finally simulated the differences in D1R expression on layer III dlPFC PV neurons between marmosets and macaques in a newly designed dlPFC microcircuit model of a visual fixation task requiring overt sustained attention and distractor resistance. Importantly, in our visual fixation model we had D1R+ Fixation I_*NEAR*_ cells. For these, we implemented the same D1R-modulated firing thresholds as for the D1R+ I_*NEAR*_ cells in the main Network 1 of the covert working memory maintenance model. This aligns the two models in the same D1R modulation framework for comparability, consistency, and generalizability. Indeed, particularly at the typical, mid D1R occupancy we observed that the macaque-D1R model of sustained attention exhibited optimal noise and distractor resistance, like the macaque-D1R model of working memory maintenance. In comparison, at the same D1R occupancy, the higher D1R expression in the Fixation I_*NEAR*_ cells, putative PV neurons, increased the distractibility of the marmoset-D1R model of sustained attention, like in the marmoset-D1R model of working memory maintenance. In addition, the visual fixation model reproduced the relative difference in the median fixation duration between marmosets and macaques in our behavioral task. Overall, our microcircuit models show that the higher D1R expression in the layer III dlPFC PV neurons of marmosets can account for their higher distractibility across tasks that rely on layer III dlPFC, through a common neurophysiological mechanism, namely the disruption of task-related persistent activity.

Interestingly, the marmoset-D1R model of sustained attention further showed higher variability in trial-by-trial fixation durations relative to the macaque-D1R model. Such higher variability is attributable to the higher excitability of D1R+ Fixation I_*NEAR*_ cells in the marmoset-D1R model (due to the higher D1R expression), which makes them more sensitive to the stochasticity of the excitatory background noise that they receive in the form of uncorrelated Poisson spike trains. In real brains, the higher D1R expression in layer III dlPFC PV neurons would make them more sensitive to fluctuations in DA release as well, which we have not simulated here but which would also affect cognitive performance. Thus, our modeling may suggest that neuroanatomical features that increase the excitatory gain of somatic PV inhibition in layer III dlPFC may make disruptions of task-related persistent activity more dependent on the stochasticity of neurophysiological processes. However, our modeling may not explain the higher variability in the mean fixation durations of the eight marmosets, relative to those of the two macaques, in our behavioral data, which was observed at the group level. Apart from the potential biases introduced by the differences in sample sizes between species, this group-level difference may instead tentatively indicate that individual differences in background noise or DA release are bigger in marmosets than macaques, but this issue is complex and outside the scope of our study.

Relatedly, for simplicity, our modeling does not simulate changes in extracellular DA concentration in dlPFC, assuming marmosets and macaques normally function at mid D1R occupancy in tasks requiring working memory maintenance or sustained attention. However, DA release varies significantly over time, across mental states, and under different physiological or environmental conditions, among other factors, in both species, and thus *in vivo* they would utilize the entire D1R occupancy scale more readily during cognitive operations, instead of primarily functioning in the mid D1R occupancy range. In that context, a reduced range of D1R occupancies that support distractor-resistant persistent activity in the marmoset-D1R model (e.g., Figure 6C for visuospatial working memory maintenance) may perhaps better explain the higher observed distractibility in marmosets.

Moreover, the modeling assumes that no systematic differences in dlPFC DA release exist *between* the species, particularly during cognitive performance. However, if dlPFC DA release is lower in marmosets than macaques, the cognitive impact of their higher D1R expression in layer III dlPFC PV neurons would be reduced, suggesting that other species differences are more likely to explain their higher distractibility. Few studies have examined the organization and function of the mesocortical DA system in marmosets, whether focusing on the cortical pattern of DA innervation, DA release, uptake, and metabolism, or extracellular DA concentration. To our knowledge, no direct comparisons exist between marmosets and macaques, especially under cognitively demanding conditions. Thus, we can only make inferences from indirect comparisons, which may be confounded by unknown and possibly substantial differences in methods or analyses between studies. Furthermore, available studies were primarily neurophysiological, and cognitive factors were not considered. Tentatively, differences appear to support a higher level of extracellular DA concentration in the cortex of marmosets^**111-113**^, which could shrink the optimal D1R occupancy range in our marmoset-D1R models (e.g., Figure 6C). In effect, the distractibility in marmosets may be even greater than what could be expected based solely on their higher D1R expression in layer III dlPFC PV neurons. Thus, the assumption that extracellular DA concentrations in dlPFC are equivalent between species, as between the macaque-D1R and the base marmoset-D1R models, may in fact be conservative. Nevertheless, direct species comparisons of extracellular DA concentration and turnover in dlPFC specifically, and ideally during working memory maintenance and sustained attention, will be needed to guide the interpretation of our results.

Otherwise, the fewer spines on the basal dendrites of pyramidal neurons, potentially leading to weaker recurrent excitation, in marmoset dlPFC^**37**^ could also significantly contribute to the higher distractibility. Accordingly, decreasing the maximum strength of Delay and Fixation Rule E→E synapses in our marmoset microcircuit models would account for this species difference. In effect, this would cap recurrent excitation to a lower level, and the D1R-mediated increase of *G*_E→E, NMDA_ would not facilitate the same amplitude of persistent activity as in the macaque-D1R model, thus decreasing the distractor resistance of the microcircuit. Additionally, marmosets have a smaller dlPFC than macaques^**34-36**^, which has been linked with limited capacity for working memory^**114**^. Furthermore, in the original spiking model of covert working memory maintenance^**64**^, decreasing the microcircuit size increased the spatial drift of persistent activity representations (away from the angular positions where stimuli appeared) due to random noise inputs. Thus, in our working memory modeling, decreasing the size of the Delay microcircuit in the marmoset model would reduce the stability of visuospatial representations against noise, and thus increase errors where the decoded position of the persistent activity trace is > +/-22.5° of the target. On the other hand, our current visual fixation microcircuit would not reproduce this effect. The latter would require modeling a spatially continuous Fixation population with neurons tuned to different eccentricities in place of the discrete, foveal Fixation population that we now have.

Importantly, numerous other neural substrates involved in working memory maintenance and sustained attention could also differ between marmosets and macaques and thus further contribute to differences in their cognitive performance. Species differences in receptors and transporters that control glutamatergic and GABAergic neurotransmission would impact the neurophysiological processes that we have modeled here. For example, as predicted by early modeling studies^**115**, **116**^, NMDARs containing the GluN2B subunit, which provides extended calcium influx^**117-119**^, are critical for persistent activity within macaque layer III dlPFC microcircuits during working memory maintenance^**120**, **121**^, and thus also possibly sustained attention. Given the increasing expression of NMDAR-GluN2B across primate phylogeny in the PFC^**122**^, differences in NMDAR-GluN2B expression in layer III dlPFC pyramidal neurons could influence discrepancies in higher cognition between marmosets and macaques. In addition, species differences in D1R expression levels or membrane localization in layer III dlPFC pyramidal neurons may also differentially affect working memory maintenance and sustained attention between marmosets and macaques. However, while transcriptomic studies are beginning to address this topic^**58**, **61**, **123**, **124**^, mRNA and protein expression can be uncorrelated^**125-127**^. Hence, a thorough, protein-level exploration of differences in the expression of D1Rs, and other molecular substrates of higher cognition, will significantly improve our understanding of the many molecular mechanisms that could produce varying cognitive performance between the species. All in all, future studies using multi-scale approaches will be crucial to capture all the molecular, cellular, microcircuit, and network differences that may contribute to cognitive and behavioral disparities across marmosets, macaques, and other primates.

Another caveat of our study is that we could not establish the contribution of species differences in peripheral factors such as oculomotor control on the differences in behavioral performance in the visual fixation task because data from a control fixation condition without distractors was not available to us. Nevertheless, we have previously shown that marmosets have a more restricted oculomotor range, which would favor them towards maintaining gaze near the position of rest for the head that is at the screen center^**57**^, an advantage for fixation which would not explain their worse performance relative to macaques. Additionally, we could not classify the outcomes of fixation breaks in our behavioral task (e.g., saccades towards distractors, anti-saccades, etc.) because the necessary high-resolution eye-tracking data from the marmosets was unavailable. However, given that distractors in the visual fixation task appeared in a random stream of unpredictable positions within each trial, we expect most behavioral responses as a result of fixation breaks to be reflexive (e.g., saccades towards the distractors or blinks) because animals would not be able to form any anticipatory strategy that would prompt saccades in opposite or unrelated directions. Furthermore, given that the layer III dlPFC microcircuitry involved in overt sustained attention blocks inappropriate, reflexive responses during the maintenance of task-related persistent activity^**77**^, we predict that the higher D1R expression in layer III dlPFC PV neurons in marmosets, which causes more prevalent disruptions of persistent activity, would specifically increase the incidence of the reflexive saccades and blinks they make, relative to macaques. Future research could test this prediction.

A final major caveat of our study is that we performed our transcriptomics, anatomical analysis, and behavioral testing in separate animal subjects. Thus, we have no within-subject evidence that species differences in D1R expression relate causally to the differences in behavioral distractibility we detected or the neurophysiological correlates we propose. Advances in genetic and molecular tools for non-human primates would allow testing such causal relationships, e.g., through cell-type-specific manipulation of D1R expression or activation in layer III dlPFC PV neurons, or otherwise of their membrane excitability more directly. Crucially, our modeling provides a strong and coherent theoretical basis for such research to be performed when possible.

The current findings suggest that DA may have a larger inhibitory effect on dlPFC microcircuits in marmosets than macaques in response to task-irrelevant environmental stimuli. The DA neurons that project to the dlPFC respond to unexpected, salient stimuli, whether they are aversive or rewarding^**41**, **128**^, and the current data suggest that moderate levels of DA release to an unexpected, salient stimulus may enhance dlPFC function in macaques, but overexcite PV neurons and thus inhibit dlPFC function in marmosets, increasing their distractibility. The higher susceptibility of cognitive processes to interference by environmental stimuli may benefit survival in smaller prey species like marmosets by lowering the threshold for predator detection and instinctive responding. This mechanism may be shared in mice, another small prey species. At higher levels of DA release during states of uncontrollable stress^**129**, **130**^, dlPFC dysfunction is also seen in macaques^**46**, **130**^ and humans^**131**^, and may contribute to symptoms of mental illness, for example, in schizophrenia^**132**^. In fact, our research suggests a testable hypothesis that the increased D1R expression observed in the dlPFC in schizophrenia^**133**^ may be partly localized to layer III PV neurons, which may be related to the working memory deficits of the disorder. Importantly, our cross-species modeling represents an initial effort in translational computational neuroscience, which may become an impactful field because *in silico* approaches can offer clear mechanistic insights and predictive power, accelerating the discovery and development of effective treatments for psychiatric disorders.

To conclude, our multi-scale study explored behavioral correlates of cognitive function, *DRD1* mRNA and D1R protein levels *in situ*, and computational modeling of molecular, cellular, and microcircuit bases of working memory maintenance and sustained attention. Our findings indicate a general mechanism by which subtle cell type-specific molecular differences in dlPFC microcircuits may significantly affect cognitive function. Specifically, we found that a difference in the expression of one receptor in one cell type, namely D1R in layer III dlPFC PV neurons, may contribute to species differences in distractibility even in a visual fixation task requiring no working memory maintenance. Notably, we empirically confirm marmosets are more distractible than macaques – a crucial step in understanding the translational validity and viability of the former as non-human primate models of cognition. Furthermore, our results suggest that natural variation in D1R expression in PV neurons in humans may contribute to variation in distractibility.

## Online Methods

### Behavior

#### Subjects

Eight marmosets (age ranging 2-7 years, 7 male and 1 female) and two macaques (age ranging 10-15 years, both male) were used. All experiments in marmosets were approved by the Institutional Animal Care and Use Committees at their respective institutions which included 5 males and 1 female at the University of Rochester, New York, and two males at the University of California, San Diego, that were part of a previous study^**1**^. All procedures in macaques were approved by the Salk Institute Institutional Animal Care and Use Committee and conformed to NIH guidelines. Fixation data from previously published studies were reanalyzed in the two macaques^**2**^ and from two male marmosets^**1**^, while new data from 6 additional marmosets trained in the same task as the original two marmosets have been contributed for this study.

#### Testing

To quantify the difference in distractibility between marmosets and macaques we compared data collected from previous studies in marmosets and macaques that used a similar basic fixation task, with no motor demands and minimal cognitive load, where the subjects ignored peripherally flashed distractors to maintain central fixation. This task is commonly used to map receptive field properties of neurons in visual areas with rhesus macaques^**2**^, and it has been shown that marmosets can also perform similar fixation tasks^**1**^. Both species underwent similar training protocols to learn how to acquire fixation on a central point and then hold it through a period in which distractors were flashed in the periphery. All subjects in the present study had been previously trained on the task during receptive field mapping for visual tasks. They were thus acclimated to the task, which precluded a learning component in the testing sessions used for the present study. Macaques and marmosets viewed a centrally presented fixation point (∼0.2-0.3° radius, at 100% contrast) on a video display at 57 cm distance while their eye position was monitored with an infra-red eye tracker. After calibration of the eye tracker, monkeys were required to maintain fixation within a 1-2° radius window around the central point. In each task trial fixation was first obtained within the fixation window and held for a minimum duration of 100 ms, after which a set of Gabor edge stimuli were flashed at peripheral positions. The flashed Gabor stimuli were 1-2° in diameter and had a spatial frequency between 1-2 cycles per degree. They were drawn at random from 8 orientations and shown at 100% Michelson contrast against a grey background. The flashed stimuli could appear at randomly selected positions ranging between +/-15 visual degrees on the vertical and horizontal but not within 2° of the fixation point. They were updated in position at each video frame at a rate of 60 Hz. Monkeys were rewarded with a drop of juice for holding fixation at increasingly longer epochs up to a maximum delay of 3 seconds. During initial training the minimum hold duration for reward was gradually increased per behavioral session for macaques until they saturated in performance for their held fixation duration. In the case of marmosets, the fixation duration was also gradually increased, but done so within a behavioral session using a 1-up 1-down staircase procedure where the duration increased after each successful hold trial and decreased upon a failed trial, and where the amount of juice reward provided was in proportion to the duration of the hold. Specifically, marmosets received more drops of liquid reward for holding longer times with a minimum reward of 2 liquid drops plus an additional drop for every 500 ms held, ranging from 2-6 total drops.

#### Data Analysis and Statistics

Data used to estimate hold durations was taken after initial training had reflected a saturation in performance. After performance saturated a minimum of 8 behavioral sessions with a minimum of 200 completed fixation trials was collected. We computed the mean fixation hold and standard deviation for each animal. Mean hold durations between macaques and marmosets were compared for a difference in median mean duration using a two-tailed Ranksum test.

#### Single nucleus RNA sequencing of DRD1 in PVALB neuron clusters of dlPFC comparing macaques and marmosets

Single nucleus cortical GABAergic neuron data were obtained and reanalyzed from a prior study by Krienen *et al*.^**3**^. Single nucleus RNA sequencing data from macaque cortical GABAergic neurons was obtained from a dataset from which only glutamatergic neurons have been previously analyzed^**4**^.

Information about the subjects used for this study and sequencing procedures can be found for marmosets^**3**^ and macaques^**4**^. Briefly, marmosets (n = 12, sex = 7F, 5M, age range 2-14 years) were cared for and housed at the Broad Institute in Cambridge, MA and all procedures were approved by the Massachusetts Institute for Technology Committee for Animal Care (CAC) guidelines. Macaques (n = 2, both female, age range 6-8 years) were cared for and housed at the University of Pittsburgh in Pittsburgh, PA and all procedures were approved by the University of Pittsburgh Institutional Animal Care and Use Committee (IACUC).

Fresh-frozen postmortem macaque dlPFC tissue containing both banks of the principal sulcus area 46 was obtained from two experimentally-naïve, female, adult (6.6 and 8.2 years of age) Rhesus macaque monkeys at the University of Pittsburgh. Animals were deeply anesthetized and perfused with artificial cerebral spinal as previously described^**4**^ in accordance with USDA and NIH guidelines and with the approval of the University of Pittsburgh IACUC.

Single nucleus RNA sequencing was performed on marmoset and macaque dlPFC tissues using the 10x Chromium 3’ v3 and v3.1 chemistry. Nuclei suspensions, library preparation, sequencing and preprocessing for marmoset is described in Krienen *et al*.^**3**^ and macaque by Datta *et al*.^**4**^. Marmoset data were aligned to genome CJ1700 from NCBI; macaque data were aligned to reference Mmul10.

Analysis and QC was performed as previously described^3, 4^. Briefly, nuclei with fewer than 500 genes or 1000 transcripts were removed from the analysis. Independent component analysis (ICA, using the fastICA package in R) was performed on all cells, using the filtered digital gene expression matrix (DGE) after normalization and variable gene selection following procedures described in Saunders *et al*.^**5**^ and Krienen *et al*.^**6**^. Louvain clustering (resolution 0.1, nearest neighbors = 25) was performed using the top 60 ICs. ICA and Louvain clustering was repeated on each of the neuron clusters, resulting in further subdivisions of the major types. Cell and gene weights on independent components (ICs) were manually inspected for evidence of cell-cell doublets or non-neurons, both of which were removed from the DGE. ICs that reflected artifactual signals or batch effects were identified and excluded, and Louvain clustering was performed on the retained ICs. Within each cluster, transcripts for each gene were summed across cells of that cluster. The resulting “metacell” counts were normalized to the total number of transcripts, then scaled to counts per 100,000 and log10 transformed.

For each species (marmoset, macaque), cortical GABAergic clusters were identified based on the expression of GABAergic neuron markers (*GAD1, GAD2, RBFOX3*), and carried forward for further analysis. Expression levels per 100k transcripts were averaged over *PVALB* clusters if there were more than one detected.

### Evolutionary transcriptomics

To plot phylogenetic trends in *DRD1* expression in PFC *PVALB* neurons, we statistically compared its expression in mice, non-human primates and humans. We first downloaded the processed single-nucleus RNA-seq data from adult primates (i.e., human, rhesus macaque and marmoset) dlPFC^**7**^. Because the dlPFC is unique to primates, we included the single-cell transcriptomics data from analogous mouse areas PL-IL-ORB (prelimbic, infralimbic and orbital regions) regions^**8**^. The *PVALB* neuron populations were extracted from both datasets.

To account for the biological variation present among donors of each species, we aggregated the UMI (unique molecular identifier) counts of *PVALB* neurons in each donor via pseudobulking strategy^**9**^. We then subset the data using the ortholog genes identified via orthogene package^**10**^ and conducted different gene expression test using edgeR package^**11**^. The ortholog genes were also leveraged to compute the library size in each pseudobulk sample followed by library size normalization and log2 transformation, using the following equation:

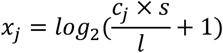

Here, the *x*_*j*_ represents the log2-normalized expression of a given gene *j, c*_*j*_ refers to the unique molecular identifier (UMI) counts for the gene *j, s* is the scaling factor (1000,000 was used here), and *l* denotes the library size.

### Evaluation of in situ protein expression of D1R in PV neurons of dlFPC

#### Subjects

Tissue for analysis of D1R protein-level expression in macaques was obtained from our extant brain bank from the Yale colony of rhesus macaques (*Macaca mulatta*). All research was approved by the Yale IACUC and was in compliance with NIH and NSF regulations. Animals were perfused with 0.1M phosphate buffer, followed by 4% paraformaldehyde and 0.05% glutaraldehyde in 0.1M PB, for concurrent studies conducted requiring fixation protocols for electron microscopy. For each subject, a craniotomy was performed, and the entire brain was removed, including a dorsolateral block containing the principal sulcus. Blocks were sectioned coronally at 60µm on a vibratome (Leica VT100S). Sections were cryoprotected through increasing concentrations of sucrose solution (10%, 20% and 30% each overnight), cooled rapidly using liquid nitrogen and stored at -80°C. To enable penetration of immunoreagents, all sections were subjected to three freeze-thaw cycles in liquid nitrogen. Sections from the dlPFC of two female young adult macaques aged 8 and 10 years were used in this study. We isolated sections with the principal sulcus and imaged in the dorsal or ventral bank of the principal sulcus, approximately midway through its anterior-posterior axis.

Tissue in marmosets was obtained from University of Western Ontario, where all experimental procedures were performed in accordance with the Canadian Council on Animal Care guidelines and approved by the University of Western Ontario Animal Use Subcommittee (AUP: 20180130) and in accordance with NSF regulations. One male and one female (both aged 5 years) young adult marmosets were perfused with 4% paraformaldehyde and 0.05% glutaraldehyde in 100mM PBS, and postfixed in 4% paraformaldehyde in 0.1M PBS for 24 hrs and blocked. Dorsolateral blocks were cut on a vibratome into 50 um sections and collected serially, and stored long-term at -20°C in cryoprotectant composed of 30% glycerol, 30% ethylene glycol in 0.1M PB. For immunohistochemistry, we sampled sections containing A46d and A46v, guided by the location of delay-related neuronal firing correlated with working memory maintenance^**12**^.

#### Antibody selection and validation

D1R labeling in macaques has been performed for decades using a knockout-validated^**13**^ antibody (Sigma rat anti-D1R, cat# D2944, AB_1840787) against the human D1R. This antibody is known for producing a patchy pattern of labeling in neurons, which has been attributed to its robust labeling in the Golgi apparatus^**14-16**^. We found that this antibody performed differently in the marmoset, producing a qualitatively distinct labeling pattern from the macaque. The labeling in the marmoset did not exhibit the characteristic patchy Golgi-like pattern well-documented in the macaque, and which we observed in our macaque tissue. The marmoset labeling pattern entirely filled the cytoplasm of labeled neurons and also labeled what appeared to be glia. This discrepancy may be due to two amino acid residue sequence differences between macaques and marmosets in the epitope region of the D1R that is bound by the Sigma antibody. We then optimized the second-best performing antibody from a knockout validation study^**13**^, namely the Alomone rabbit anti-D1R (cat# ADR-001, AB_2039826), which has been used successfully in macaque tissue for immunofluorescence^**17**^, and produced satisfactory and comparable labeling in our macaque and marmoset tissue, although without the characteristic Golgi-like labeling pattern observed by the Sigma antibody. The Alomone antibody was generated against an epitope on the intracellular C-terminus of the rat D1R (amino acid residues 372-385). The amino acid residues in the epitope sequence are conserved in marmosets and macaques. We used the Alomone D1R blocking peptide (cat# BLP-DR001) as a negative preadsorption control, described more fully below.

#### Immunohistochemistry

Both species were processed together using the same solutions to ensure identical treatment. Free-floating sections were removed from long-term storage and washed in phosphate buffer (0.1M PB, 4×15 min), followed by an incubation for 1 hr in glycine (50 mM in 0.1M PB, room temperature, Sigma, cat# G-7126) to bind free aldehydes, PB washes (4×15 min), and Avidin Biotin blocking (Vector, cat# SP-2001) to block any endogenous biotin. Following more PB washes (4×15 min), the tissue was incubated for 1 hr at room temperature in a blocking solution of 5% bovine serum albumin (AmericanBio, cat# 9048-46-8), 10% normal goat serum (Jackson ImmunoResearch, cat# 005-00-001), and 2% Triton X-100 (Electron Microscopy Sciences, cat# 22140) in 0.1M PB. Tissue was then incubated for 72 hrs at 4°C in PB with rabbit anti-D1R (1:100, Alomone, cat# ADR-001), and guinea pig anti-PV (1:2000, Swant, cat# GP72, AB_2665495). Tissue was then washed in PB (4×15 min) and incubated for 3 hrs at room temperature in PB with Alexa Fluor-488 goat anti-rabbit (1:100, Invitrogen, cat# A11008) and Alexa Fluor-568 goat anti-guinea pig (1:100, Invitrogen, cat# A11075). Sections were then washed with PB (4×15 min), and mounted onto slides using Prolong Gold Antifade (Invitrogen, cat# P36930), dried, coverslipped, and sealed with fast-drying nail polish. A control piece of tissue was treated as above, except for the primary antibody incubation step, which additionally included the Alomone D1R antigen blocking peptide (cat# BLP-DR001) at 10x the D1R antibody concentration (pre-incubated for 1 hr). Upon imaging, no material labeling was observed in the D1R channel in the control section (Figure S3).

We also performed additional immunofluorescence labeling, depicted in Figure S1. D1R and PV labeling procedures were performed as above with the addition of microtubule-associated protein 2 (MAP2) and the nuclear Hoechst stain. The primary antibody incubation included the addition of a chicken anti-MAP2 antibody^**18**^ (Abcam, cat# AB5392, 1:1000, AB_2138153), and the secondary antibody incubation included goat anti-chicken Alexa Fluor-647 (1:100, Invitrogen cat# A21449). The Hoechst nuclear stain was included at the end at 1:10,000 (Fisher Scientific, cat# H3570).

#### Sampling and Imaging

In layer IIIb, 5-10 sampling sites were imaged per section, sometimes capturing almost all of the marmoset 46d or 46v layer III in section, and in macaque utilizing every other field of view traversing along deep layer III parallel to the pia. Layers were determined at a lower magnification objective using the pattern of pyramidal D1R labeling, and PV labeling, which is specific to laminar compartments^**19**^. Confocal images for marmosets and macaques were acquired during the same session using a Zeiss LSM880 Airyscan with the Plan-Apochromat 20x/0.8 M27 or C-Apochromat 40x/1.2W Korr FCS M27 objectives. Z-stacks at various z-steps were obtained under sequential excitation at 488 nm, and 561 nm. After initially surveying sections from both species, imaging parameters (laser power, gain settings, pinhole, etc.) were set at the start so that images acquired across species were subject to equivalent imaging parameters. Parameters were optimized for focal planes near the top and bottom of the tissue section, as D1R antibody penetration was attenuated in the middle depths of the sections for both species, and to avoid pixel saturation. Images were deconvolved using Huygens Professional version 22.04 (Scientific Volume Imaging, The Netherlands) using as uniform parameters as possible (theoretical PSF, classic MLE algorithm, SNR set at ∼15, background set at 0, and acuity optimized for smoothness). We used Fiji^**20**^ to further analyze all deconvolved images. The D1R signal was optimal in a window of ∼15 µm from the top or bottom of the section. Two single focal planes from each stack were extracted, one each from the optimal sectors at the top or bottom of the section, while ensuring that no neurons were present in both and avoiding the absolute top or bottom due to the presence of surface artifacts. Extracted focal planes were used for two independent analyses, discussed sequentially below. We sampled from 4 sections per subject. In total, we analyzed ∼100 images, encompassing ∼400-500 PV neurons per subject.

#### Analysis of D1R expression in PV processes

To perform the colocalization analysis, we used the EZColocalization plugin^**18**, **21**^. For each extracted focal plane, we performed particle segmentation to isolate PV neurons and dendrites, utilizing either the Renyi or Otsu thresholding methods, producing sometimes over a thousand particles per image, including both dendritic and somatic segments. An areal filter was applied so particles smaller than 0.55 µm^2^ were excluded. Segmented particles were analyzed manually to classify into two categories, i) PV neuron somata and/or proximal dendritic segments, which were sometimes continuous; or ii) non-proximal PV dendrites. Any quantal isolated signals that could not be traced to a continuous dendrite were discarded and proximal dendrites were identified as being readily traced back to a PV+ neuron soma. The areal segmentation filter and manual identification helped exclude PV+ thalamocortical axons that may have been present within the section. For each focal plane, PV+ particles were then analyzed for colocalization using the Costes’ thresholding method and Manders’ Overlap Coefficient (MOC). Briefly, the Costes’ thresholding identifies pixels that qualify as colocalized for each image, and the MOC roughly computes the proportion of colocalized pixels among total pixels contained in a PV particle. We ran a separate Costes’ and MOC analysis on a subset of manually segmented PV particles and received consistent results with the automatically segmented set.

Then, we performed a separate independent analysis of D1R labeling in manually traced PV neuron somata. This is because the automated segmentation often produced multiple segments per PV soma, and included dendritic PV segments present in the image. For each image, we isolated the D1R channel and manually traced the 3-6 most strongly D1R labeled somata in the image, which were typically pyramidal-like in morphology. We then sampled 10-20 segments of the immunonegative “neuropil”. In the isolated D1R channel, we measured the mean grey value (MGV) for the strongly labeled D1R somata traces and the sampled neuropil traces. We then used the minimum MGV across the “strong” D1R traces, and the average MGV across D1R “neuropil” traces to compute two bins of intermediate D1R labeling between the average MGV in the neuropil and the minimum MGV from the strongly labeled D1R traces. In this way, we were able to normalize for the illumination level of each image. Then, in the isolated PV channel of the same image, we traced the outlines of the PV somata, measured the D1R MGV for each traced PV somata, and classified the PV somata as having a D1R MGV i) below the average MGV in the neuropil, deemed “D1R negative”, ii) in two intermediate bins, “D1 weak” or “D1 moderate”, or iii) above the minimum MGV observed among the “strongly” labeled D1R somata. This strategy has been schematized in Figure S2. We analyzed a total of 1,821 PV neuronal somata across all sampled focal planes (Marm1: n = 512 PV neurons; Marm2, n = 501; Mac2, n = 403; Mac1, n = 405).

Brightness and contrast adjustments were made for qualitative figures using Fiji or Adobe Photoshop CS5 Extended (version 12.0.4×64, Adobe Systems Incorporated), though all quantitative analysis was performed on untouched images straight after deconvolution. Figures were prepared in Adobe Illustrator CC.2020.01 (Adobe Systems Incorporated).

#### Statistical Analysis

Statistical analysis was performed using Prism 9 (Graphpad Software, LLC). To remove independence violations, we analyzed the data using the mean per subject. We checked the assumptions for parametric tests, and used a t-test to test the mean MOC across species, and a two-way ANOVA with post-hoc Sidak’s corrected pairwise comparisons to test the effects of species and effect of proximity to soma, and interaction between the two. Means are reported with standard deviation, when appropriate, as indicated in the text and plots.

### Computational Modeling

#### Single-cell model

We modelled all cells in the spiking network models as single-compartment leaky integrate-and-fire units characterised by the following intrinsic parameters: total membrane capacitance (*C*_*m*_), total leak conductance (*g*_*L*_), leak reversal potential (*E*_*L*_), the firing threshold potential (*V*_*th*_), the reset potential (*V*_*res*_), and the refractory time (*τ*_*ref*_)^**22**^. For E and I cells, the parameter values used in simulations are listed in Table 1^**23**, **24**^. The change in the subthreshold membrane voltage *V* at a given time point *t* of each postsynaptic cell *i*, denoted as *V*_*i*_(*t*), is given by:

**Table 1.**
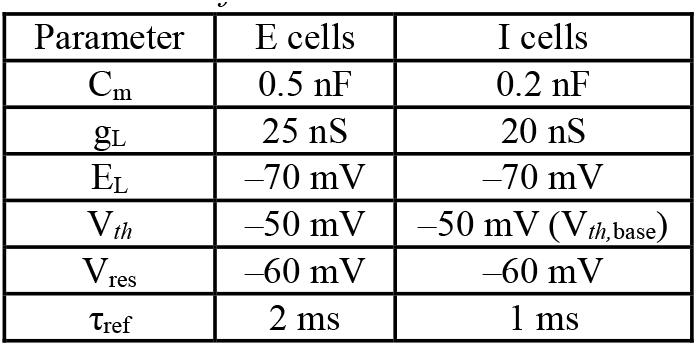
Parameters of E and I cells.

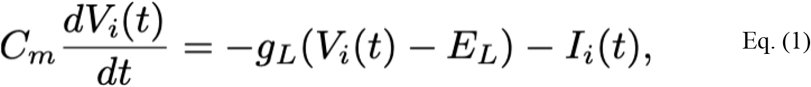

where *I*_*i*_(*t*) is the tota; input current to the cell.

#### Receptor dynamics

Cells receive recurrent excitatory inputs through α-amino-3-hydroxy-5-methyl-4-isoxazolepropionic acid receptor (AMPAR)- and N-methyl-D-aspartate receptor (NMDAR)-mediated synaptic current transmissions and inhibitory inputs through gamma-aminobutyric acid type A receptor (GABA_*A*_R)-mediated synaptic current transmissions. The total input current to a given postsynaptic cell, *I*_*i*_(*t*), is:

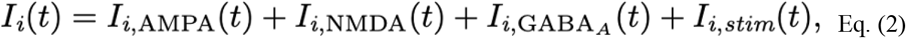

where *I*_*i,stim*_ designates the external stimulus input to the cell (see Eq. 15).

We modelled the dynamics of the receptor-specific synaptic input currents for each postsynaptic cell at a given time point, denoted as *I*_*i,r*_(*t*), based on Wang^**24**^, as follows:

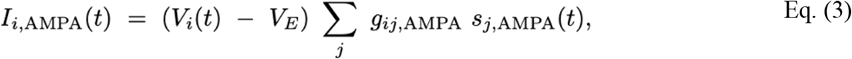

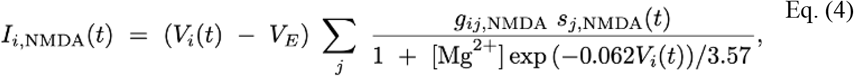

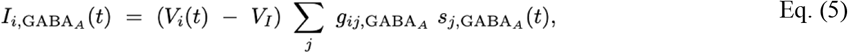

where *g*_*ij,r*_, *r* ∈ {*AMPA, NMDA, GABA*_*A*_}, is the receptor-specific conductance at the connection between presynaptic and postsynaptic cells *j* and *i. s*_*j,r*_(*t*) is the contribution to the receptor-specific synaptic gating variable by a given presynaptic cell at a given time point. *V*_*E*_ and *V*_*I*_ are the reversal potentials for the excitatory AMPAR and NMDAR channels and the inhibitory GABA_*A*_R channels (Table 2). *V*_*i*_(*t*) is the membrane potential at a given time point. The NMDAR-mediated excitatory postsynaptic currents exhibit voltage dependence governed by the extracellular concentration of magnesium, [Mg^2+^] (Table 2)^**25**^.

**Table 2.** Parameters of NMDAR, AMPAR, and GABA_A_R dynamics.

| Receptor | $V$ | $\tau$ | [Mg<sup>2+</sup>] | $\tau_x$ | $\alpha_s$ |
| --- | --- | --- | --- | --- | --- |
| NMDAR | 0 mV | 100 ms | 1 mM | 2 ms | 500 Hz |
| AMPA | 0 mV | 2 ms |  |  |  |
| GABA<sub>A</sub>R | 70 mV | 10 ms |  |  |  |

We modelled the AMPAR and GABA_*A*_R synaptic gating variables as instantaneous jumps of magnitude 1 when a presynaptic spike occurs, followed by a fast exponential decay with time constants *τ*_AMPA_^**26**, **27**^ and 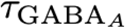^***28***, ***29***^ (Table 2). The gating variables for AMPAR and GABAR channels are, thus, given by:

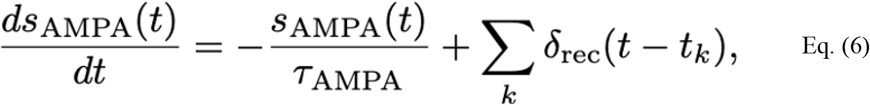

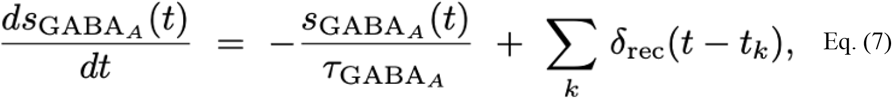

whereas the gating variable for NMDAR channels is given by:

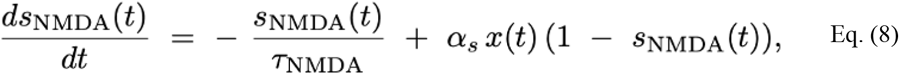

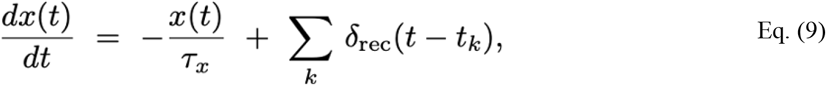

where *x* is an intermediate gating variable proportional to the glutamate concentration in the synapse, *τ*_NMDA_ is the slow decay time constant of NMDAR-mediated postsynaptic currents, *τ*_*x*_ is the rise time constant of NMDAR channels, and *α*_*s*_ is the presynaptic firing frequency at which NMDAR channels saturate and become insensitive to further frequency increases (Table 2). Across equations, *δ*_rec_ denotes the Dirac delta functions that represent the arrival of the spikes from the presynaptic, recurrent sources at times *t*_*k*_.

#### Visuospatial working memory microcircuit architecture

The three main spiking neural networks comprise 2048 near feature-tuned excitatory (E) Delay cells, 256 near feature-tuned inhibitory (I_*NEAR*_) Delay cells, and 256 opposite feature-tuned inhibitory (I_*OPP*_) Delay cells. Alternatively, in addition to 2048 E cells, the I cells in the I_*NEAR*_-only network represent 512 I_*NEAR*_ cells, while in the I_*OPP*_-only network, they represent 512 I_*OPP*_ cells. Each Delay cell type uniformly covers all angles θ on a ring in the functional feature space from 0° to 360° (Figure 4A1). E cells are indexed by the angular position of the visual stimuli to which they respond preferentially with the highest strength, while I cells conform to the indexing of the E cells from which they receive the strongest excitation.

The visuospatial working memory model features E→E (Figure 4A2), E→I, I→E (Figure 4A3-4), and I→I synaptic connections between the Delay cells. Excluding autapses, we assume all-to-all connectivity between cells for each connection type. E cells innervate with their strongest glutamatergic projections the cells of all types that occupy the nearest angular positions. I_*NEAR*_ cells recapitulate the targeting of E cells, while for I_*OPP*_ cells, the targeting is rotated by 180°. Thus, I_*NEAR*_ cells innervate with their strongest GABAergic projections the cells of all types that also occupy the nearest angular position, while I_*OPP*_ cells – the cells of all types positioned 180° away on the ring.

#### Visual fixation microcircuit architecture

The spiking neural network comprises 2048 near feature-tuned excitatory Cue (E^*CU*E^) cells, 512 near feature-tuned inhibitory 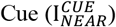 cells, 256 co-tuned, homogenously interconnected regular-spiking Fixation Rule E (E^*FIX*^) cells, 32 near feature-tuned inhibitory Fixation 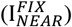 cells, and 32 opposite feature-tuned inhibitory Fixation 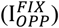 cells. The Cue cells (E^*CU*E^ and 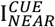) uniformly cover all angles θ on a ring in the functional feature space from 0° to 360° (Figure 7A1). Like the Delay E cells in the visuospatial working memory model, the E^*CU*E^ cells here are indexed by the angular position of the visual stimuli to which they respond preferentially with the highest strength, while 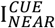 cells conform to the indexing of the E^*CU*E^ cells from which they receive the strongest excitation. The Fixation cells (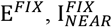, and 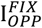) are positioned in the center of the functional feature space of the model.

The visual fixation model features E^*CU*E^→E^*CU*E^ (connectivity profile similar to that in Figure 4A2),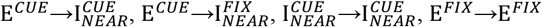 (Figure 7A2), 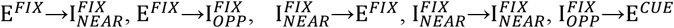synaptic connections. Notably, the 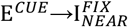 and 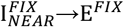 synaptic connections form a local feedforward inhibitory motif (Figure 7A3), while the 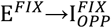 and 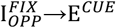 (Figure 7A4) synaptic connections form a local top-down inhibitory motif. Excluding autapses, we assume all-to-all connectivity between cells for each connection type. E^*CU*E^ cells innervate with their strongest glutamatergic projections the other E^*CU*E^ and the 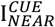 cells that occupy the nearest angular positions, as well as all 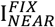 cells. E^*FIX*^ cells innervate with glutamatergic projections of the same respective strength (for a given connection type, but different strengths for different connection types) the other E^*FIX*^ cells, the 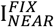 cells, and the 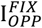 cells. 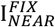 cells innervate with GABAergic projections of the same respective strength (again, for a given connection type, but different strengths for different connection types) the other 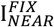 cells, the E^*FIX*^ cells, and the 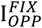 cells.

#### Synaptic connectivity

Between the Delay cells and between the Cue cells in, respectively, the visuospatial working memory model and the visual fixation model, the receptor conductance, denoted as *g*_*ij,r*_ at a connection between the presynaptic and postsynaptic cells *j* and *i* is given by:

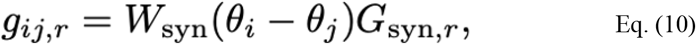

where *G*_syn,*r*_ is the mean receptor-specific conductance at the given synaptic connection type, weighted by a Gaussian distribution of the difference between the angular positions *θ*_*i*_ and *θ*_*j*_ of the preferred stimuli of the postsynaptic and presynaptic cells:

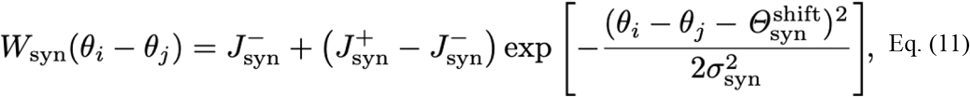

where 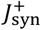 and 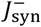 are the upper and lower limits of the Gaussian distribution function and represent the maximum and minimum synaptic connection strengths. *σ*_syn_ is the standard deviation of the Gaussian distribution function and represents the synaptic connection width. 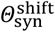 is the angular shift in the position of the set of cells targeted with the maximum connection strength relative to the position of the presynaptic cell. 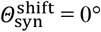 for synapses where projections originate from E or I_*NEAR*_ cells (for both the Delay cells and the Cue cells in the respective models), such that the Gaussian is centred at *θ*_*i*_ – *θ*_*j*_ = 0°. 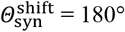 for synapses where projections originate from Delay I_*OPP*_ cells, such that the Gaussian is centred at *θ*_*i*_ – *θ*_*j*_ = 180°. 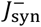 is given by the normalisation of *W*_syn_(*θ*_*i*_ – *θ*_*j*_):

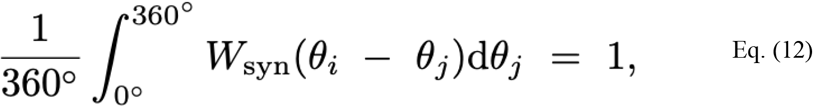

such that:

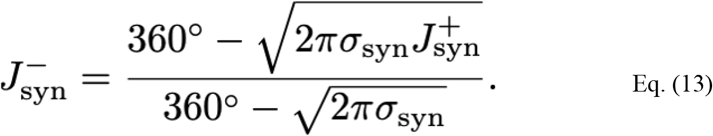

The 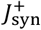 and *σ*_syn_ parameters for each synaptic connection type between the Delay cells and the Cue cells in, respectively, the visuospatial working memory model and the visual fixation model are shown in Table 3. Importantly, in the visual fixation model, the strength of all synaptic connections involving presynaptic or postsynaptic Fixation cells does not depend on *W*_syn_(*θ*_*i*_ – *θ*_*j*_), i.e. *W*_syn_(*θ*_*i*_ – *θ*_*j*_) = 1.

**Table 3.** Widths (adopted from Cano-Colino et al.^**51**^) and maximum strengths of each synaptic connection type between the Delay and Cue functional cell types in, respectively, the visuospatial working memory and visual fixation models.

| Synaptic Connection Type | Functional Cell Type | $\sigma$ | $J^+$ |
| --- | --- | --- | --- |
| E→E | Delay & Cue | 14.4° | 2.1 |
| E→I | Delay & Cue | 14.4° | 1.6 |
| I→E | Delay & Cue | 14.4° | 1.6 |
| I→I | Delay & Cue | 14.4° | 2.1 |

**Table 4.**
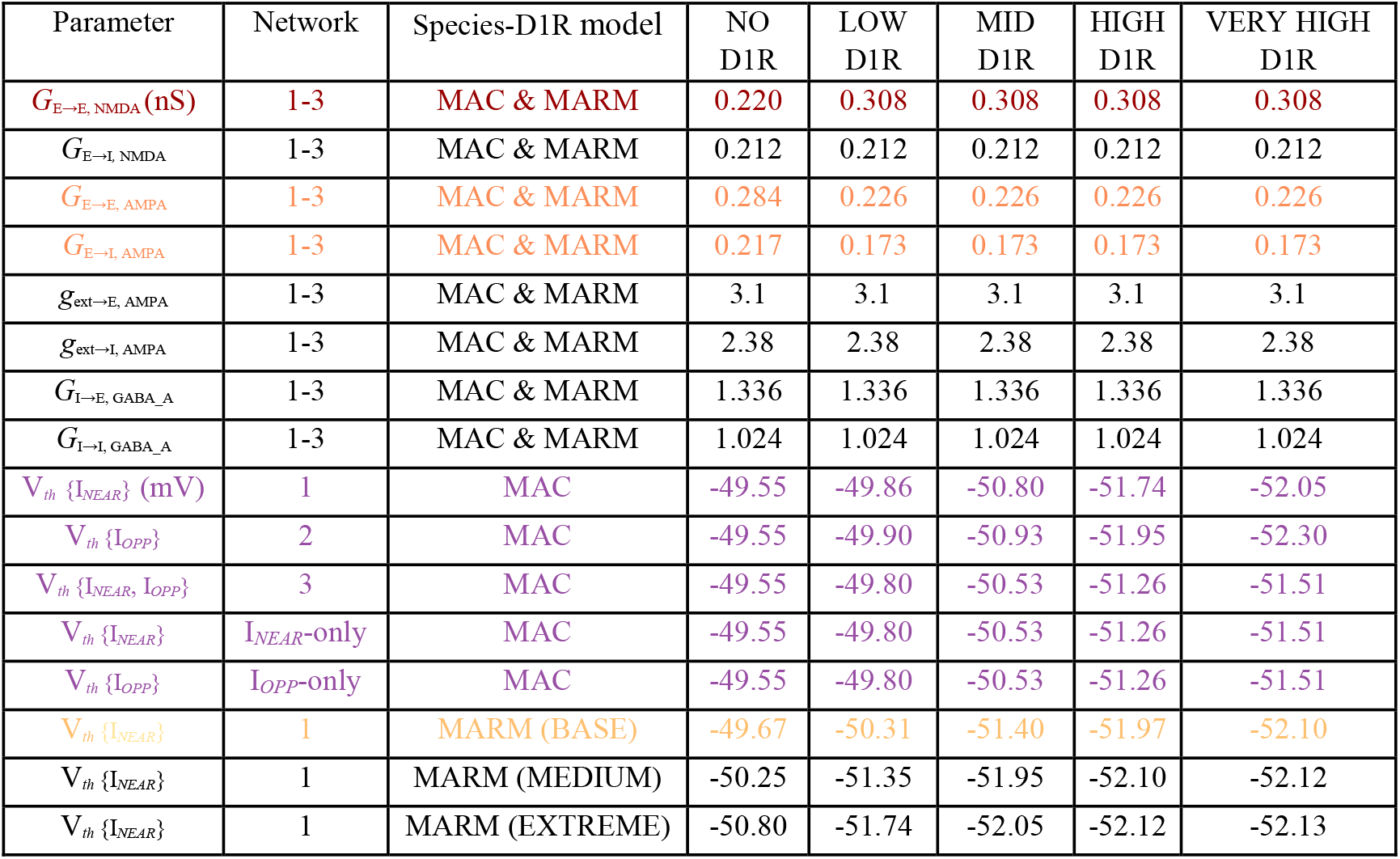
Synapse-specific receptor conductances and I cell firing thresholds for each network, species-D1R model, and D1R occupancy in the visuospatial working memory microcircuit model. Colour-coding is according to the D1R modulations represented in Figure 4C1-2.

To simulate background noise (e.g., from other cortical areas), we applied 1000 random, exclusively AMPAR-mediated, external excitatory inputs to each cell as uncorrelated presynaptic Poisson spike trains. Each train had a firing rate of 1.8 Hz, with conductances of *g*_ext→E, AMPA_ and *g*_ext→I, AMPA_ on all E and I cells in the visuospatial working memory model and the visual fixation model (Tables 4 and 5).

**Table 5.** Synapse-specific receptor conductances and 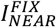 cell firing thresholds for each species-D1R model and main level of D1R occupancy on 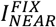 cells in the visual fixation microcircuit model. Colour-coding is according to the D1R modulations represented in Figure 4C2.

| Parameter | Species-D1R model | NO D1R | LOW D1R | MID D1R | HIGH D1R | VERY HIGH D1R |
| --- | --- | --- | --- | --- | --- | --- |
| $G_{E^{CUE} \rightarrow E^{CUE}, NMDA}$ | MAC & MARM | 0.220 | 0.220 | 0.220 | 0.220 | 0.220 |
| $G_{E^{CUE} \rightarrow I_{NEAR}^{CUE}, NMDA}$ | MAC & MARM | 0.212 | 0.212 | 0.212 | 0.212 | 0.212 |
| $G_{E^{CUE} \rightarrow E^{CUE}, AMPA}$ | MAC & MARM | 0.284 | 0.284 | 0.284 | 0.284 | 0.284 |
| $G_{E^{CUE} \rightarrow I_{NEAR}^{CUE, FIX}, AMPA}$ | MAC & MARM | 0.217 | 0.217 | 0.217 | 0.217 | 0.217 |
| $G_{E^{FIX} \rightarrow E^{FIX}, NMDA}$ | MAC & MARM | 0.308 | 0.308 | 0.308 | 0.308 | 0.308 |
| $G_{E^{FIX} \rightarrow I_{NEAR, OPP}^{FIX}, NMDA}$ | MAC & MARM | 0.212 | 0.212 | 0.212 | 0.212 | 0.212 |
| $G_{E^{FIX} \rightarrow E^{FIX}, AMPA}$ | MAC & MARM | 0.284 | 0.284 | 0.284 | 0.284 | 0.284 |
| $G_{E^{FIX} \rightarrow I_{NEAR, OPP}^{FIX}, AMPA}$ | MAC & MARM | 0.217 | 0.217 | 0.217 | 0.217 | 0.217 |
| $g_{ext \rightarrow E^{CUE, FIX}, AMPA}$ | MAC & MARM | 3.1 | 3.1 | 3.1 | 3.1 | 3.1 |
| $g_{ext \rightarrow I_{NEAR, OPP}^{CUE, FIX}, AMPA}$ | MAC & MARM | 2.38 | 2.38 | 2.38 | 2.38 | 2.38 |
| $G_{I_{NEAR}^{CUE} \rightarrow E^{CUE}, GABA_A}$ | MAC & MARM | 1.503 | 1.503 | 1.503 | 1.503 | 1.503 |
| $G_{I_{NEAR}^{FIX} \rightarrow E^{FIX}, GABA_A}$ | MAC & MARM | 1.703 | 1.703 | 1.703 | 1.703 | 1.703 |
| $G_{I_{OPP}^{FIX} \rightarrow E^{CUE}, GABA_A}$ | MAC & MARM | 0.334 | 0.334 | 0.334 | 0.334 | 0.334 |
| $G_{I_{NEAR}^{CUE} \rightarrow I_{NEAR}^{CUE}, GABA_A}$ | MAC & MARM | 1.024 | 1.024 | 1.024 | 1.024 | 1.024 |
| $G_{I_{NEAR}^{FIX} \rightarrow I_{NEAR, OPP}^{FIX}, GABA_A}$ | MAC & MARM | 1.024 | 1.024 | 1.024 | 1.024 | 1.024 |
| $V_{th} \{I_{NEAR}^{FIX}\}$ | MAC | -49.55 | -49.86 | -50.80 | -51.74 | -52.05 |
| $V_{th} \{I_{NEAR}^{FIX}\}$ | MARM | -49.67 | -50.31 | -51.40 | -51.97 | -52.10 |

#### D1R modulation in visuospatial working memory model

We modelled the effects of D1R occupancy on the NMDAR conductance at E→E synapses (*G*_E→E, NMDA_), the AMPAR conductances at E→E and E→I synapses (*G*_E→E, AMPA_ and *G*_E→I, AMPA_), and the firing thresholds (V_*th*_) of D1R+ I cells in the respective networks, as illustrated in Figure 4C. Notably, studying any other more complex D1R expression patterns within the different cell types was prohibited by the combinatorial explosion of possible patterns. Table 4 shows the NMDAR, AMPAR, and GABA_*A*_R conductances and the decreases in the V_*th*_ of D1R+ I cells across networks and species-D1R models for each main D1R occupancy level.

From no to low D1R occupancy, specifically at *x* = 0.125, *G*_E→E, NMDA_ increased in a step-like manner by 40%, mediated by postsynaptic D1Rs^**30**, **31**^, to facilitate a transition from a transient to a persistent firing activity regime, while *G*_E→E, AMPA_ and *G*_E→I, AMPA_ decreased, also step-like, by 20%, which aimed to capture a reduction of glutamate release putatively mediated by presynaptic D1Rs^**30**, **32**^, to reduce the impact of noise-evoked activations of Delay E cells on excitatory transmission to the other Delay E cells and the Delay I cells. For mid, high, and very high D1R occupancy, *G*_E→E, NMDA_, *G*_E→E, AMPA_, and *G*_E→I, AMPA_ were unchanged. *G*_E→I, NMDA_ was not modulated across D1R occupancy levels because D1Rs and NMDARs do not interact at E→I synapses in layer III of primate (macaque) dlPFC^**32**^. We adopted Compte *et al*.’s^**33**^ inhibitory GABA_*A*_R conductances, which were unchanged across D1R occupancy levels because D1R stimulation does not modify action-potential independent GABA release from the presynaptic axon terminals of fast-spiking, putative PV neurons in layer III of primate (macaque) dlPFC^**34**^. Instead, based on Kröner *et al*.’s^**34**^ finding that a V_*th*_ decrease accompanies the increase in intrinsic excitability of these inhibitory neurons with D1R stimulation, we decreased the V_*th*_ of D1R+ I cells across D1R occupancy levels according to network- and species-D1R model-specific, shifted, and scaled sigmoid functions:

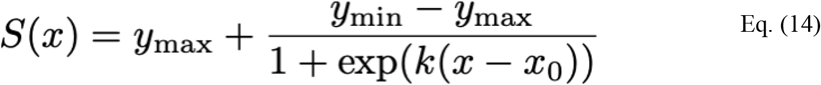

The D1R expression in PV neurons, measured by the MOC in the automated classification analysis, determined the midpoint parameters, *x*_*0*_, of the sigmoid for each species-D1R model. Specifically, we set *x*_*0*_ = 0.50 for the macaque-D1R model as the reference and *x*_*0*_ = 0.36 for the marmoset-D1R model (base case #1). However, since MOC-measured D1R expression may not directly translate to D1R stimulation, we further explored two additional cases for the marmoset-D1R model. Case #2 is characterised by a medium midpoint shift of 0.375 to *x*_*0*_ = 0.125, while case #3 – by an extreme midpoint shift of 0.50 to *x*_*0*_ =

0. Notably, leftward shifts smaller than the base case would have resulted in smaller-to-no differences in distractor resistance at the typical, mid D1R occupancy and hence were not explored. Heuristically, we choose a slope *k* = -7 to differentiate the idealised effects of the I cell thresholds on persistent activity for the five main D1R occupancy levels. *x* represents the normalised range of D1R occupancy from 0 to 1. Crucially, the same D1R occupancy signifies a greater total number of DA-bound D1Rs, and thus greater D1R modulation, in the marmoset-D1R model compared to the macaque-D1R model. Thresholds decreased for D1R+ I_*NEAR*_ cells in Network 1, D1R+ I_*OPP*_ cells in Network 2, and both D1R+ I cell types in Network 3 (Figure 4C2).

To identify the reasonable parameter sets that reproduce the inverted-U response in the macaque-D1R model, we first conducted grid searches over (1) *G*_E→E, NMDA_, (2) *G*_E→E, AMPA_ and *G*_E→I, AMPA_, and (3) V_*th*_ for the D1R+ (i) I_*NEAR*_, (ii) I_*OPP*_, and (iii) I_*NEAR*_ and I_*OPP*_ cell types in, respectively, Networks 1, 2, and 3. For comparability, the *G*_E→E, NMDA_ for no D1R occupancy was approximately the highest *G*_E→E, NMDA_ not supporting persistent activity in any network. Also, we chose the minimum threshold (V_*th*,min_) for each network to fall approximately just beyond the bifurcation point up to which persistent activity could reliably form during the delay period. Notably, we selected the V_*th*_ at mid D1R occupancy (*x*_*0*_ = 0.50) for each network to be the centre of the V_*th*_ parameter space for the spatially precise, noise- and distractor-resistant persistent activity regime, based on our assumption that this is the typical D1R stimulation level at which macaques function optimally under normal conditions. The baseline threshold (V_*th*,base_) was the value located at an equal parameter distance from the V_*th*_ at mid D1R occupancy, as compared to the distance between the latter and V_*th*,min_ due to the symmetry of the sigmoid around the midpoint. Thus, interestingly, V_*th*,base_ was -49.55 mV across networks. Then, to fit the V_*th*_ sigmoid (*x*_*0*_ = 0.50, *k* = -7) for each network to the values identified in the grid searches, we estimated the upper and lower asymptotic limits, *y*_max_ and *y*_min_, to ensure that at *x* = 0, V_*th*,base_ = -49.55 mV and at *x* = 1, V_*th*,min_ equalled a value that was different across networks (Table 4). The respective *y*_max_ and *y*_min_ are given in Table 6. Overall, for the macaque-D1R model, in Network 1, I_*NEAR*_ thresholds decreased from V_*th*,base_ for no D1R occupancy until a V_*th*,min_ of –52.05 mV for very high D1R occupancy. In Network 2, V_*th*,min_ was -52.30 mV because the model sustained persistent activity over a larger V_*th*_ range than Network 1. In Network 3, V_*th*,min_ was – 51.50 mV because the higher net inhibition, due to decreasing the V_*th*_ of twice as many I cells (512, vs. 256 in Networks 1 and 2), shortens the V_*th*_ range that supports persistent activity. Consequently, the extent of V_*th*_ decreases across D1R occupancy levels in Network 3 was smaller than in Network 1, which in turn was smaller than in Network 2. Finally, we shifted the midpoint for Network 1’s V_*th*_ sigmoid for the three marmoset-D1R model cases, while keeping all other parameters the same.

**Table 6.** Upper and lower asymptotic limits, y_max_ and y_min_, of the sigmoid functions for each model (and the networks of the visuospatial working memory model).

| Model | Visuospatial Working Memory |  |  |  |  | Visual Fixation |
| --- | --- | --- | --- | --- | --- | --- |
| Network | 1 | 2 | 3 | $I_{NEAR}$-only | $I_{OPP}$-only | |
| $y_{max}$ | -49.47 | -49.46 | -49.49 | -49.49 | -49.49 | -49.47 |
| $y_{min}$ | -52.13 | -52.39 | -51.57 | -51.57 | -51.57 | -52.13 |

We also fitted the D1R modulations of *G*_E→E, AMPA_ and *G*_E→I, AMPA_, together with the other D1R effects, to further neurophysiologically constrain network dynamics. Notably, presynaptic D1R-mediated reduction of fast non-NMDAR-mediated excitatory postsynaptic currents (EPSCs) (typically simulated by reducing *G*_AMPA_ in conductance-based models like ours) has been shown for E→I synapses in macaque layer III dlPFC^**32**^. However, to our knowledge, a similar explicit effect has not been reported for E→E synapses in primate layer III dlPFC. Furthermore, species, region, and layer differences in the D1R modulation of non-NMDAR-mediated EPSCs appear prevalent at E-E synapses (see Table 1 in the following works^**35**, **36**^), in contrast to the well-established D1R-mediated increase in NMDAR-mediated EPSCs. The most relevant study on this issue showed a reduction in AMPAR-mediated EPSCs in macaque layer III dlPFC^**37**^, but that effect was mediated by D1R and D2R co-activation and seemed to be specific to E→E synapses where the excitatory projection was long-range, with origins outside the microcircuit. Hence, we modelled the reduction in *G*_E→I, AMPA_ based on data from rat layer V mPFC instead^**30**^. Nevertheless, a computational study^**38**^ showed that neither decreases nor increases of both *G*_E→E, AMPA_ and *G*_E→I, AMPA_ (concurrently) significantly or qualitatively changed the inverted-U relationship between working memory precision (i.e., the spatial tuning of the population activity of excitatory Delay cells) and D1R stimulation, possibly because AMPARs contribute little to persistent activity^**24**, **39**^ (but see^**40**^). Thus, ultimately, any different direction of the D1R effect on *G*_E→E, AMPA_ and/or *G*_E→I, AMPA_ should not change our conclusions.

#### D1R modulation in visual fixation model

We simulated the effects of D1R occupancy only on the V_*th*_ of 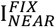 cells. Importantly, we used the D1R-modulated V_*th*_ of Delay I_*NEAR*_ cells from Network 1 of the visuospatial working memory model for the D1R-modulated V_*th*_ of 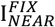 cells across D1R occupancy levels to help compare and interpret the two microcircuit models in the same framework. Notably, while E^*FIX*^ cells exhibit persistent activity during overt sustained attention^**41-46**^, there is no evidence currently that they express D1Rs, and hence we did not simulate the effects of D1R occupancy on 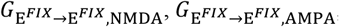, and 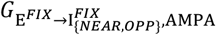. Cholinergic stimulation of nicotinic alpha2beta4 receptors does modulate the persistent activity of E^*FIX*^ cells^**41**^, but implementing this is outside the scope of our study. Regardless, modelling the transition from a distractible to an erroneous persistent activity regime (in the increasing phase of the inverted U relationship between neuromodulation and persistent activity) via modulation of 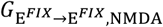 is not consequential for our goals. For this reason, we used the value of the 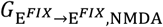 parameter from the low-to-very high D1R occupancy levels in the visuospatial working memory model for the typical persistent activity regime of E^*FIX*^ cells because that is sufficient to support persistent activity but further research would need to confirm whether stimulation of D1Rs or other receptors modulates 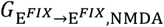. Additionally, we used the values of 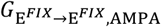 and 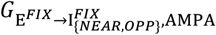 from the no D1R occupancy level in the visuospatial working memory model for the typical persistent activity regime of the visual fixation model to differentiate it from the typical persistent activity regime of visuospatial working memory model and reflect that overt sustained attention is by definition externally-driven, and hence more dependent on fast AMPAR dynamics than covert working memory maintenance. However, again, future research should test these assumptions. Table 5 shows the NMDAR, AMPAR, and GABA_*A*_R conductances and the decreases in the V_*th*_ of 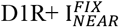 across networks and species-D1R models for each main D1R occupancy level.

#### Simulation protocol for visuospatial working memory model

We simulated five-second trials of a visuospatial working memory maintenance task mimicking Funahashi *et al*.’s protocol^**47**^, but without the fixation demands or explicit saccadic responses. For a given trial, at a time point of *T*_*targ*_ = 400 ms, for a duration of Δ*t* = 250 ms, we applied a target stimulus as a current injection of strength *μ* = 0.20* nA or 0.25** nA onto each Delay E cell within an arc with a width of *σ* = 30° centred at an angle of *Θ*_*targ*_ = 0° on the ring. A 2050-ms target delay period followed. To test the distractor resistance of the microcircuit, we then applied a distractor stimulus at *T*_*dist*_ = 2700 ms, which was always of the same strength, width, and duration as the target stimulus but centred at *Θ*_*dist*_ = 180° (in Figure 5 and for the analysis of the inverted-U response across stimulus strengths in Figure S4 because at this position studies^**33**, **48**^ suggest distractor resistance would be most reliable under optimal conditions, maximizing our chances of capturing the peak of the inverted-U) or another position (for the analysis of the inverted-U response across distractor positions in Figure S5). The distractor delay period was also 2050 ms until the end of the trial. Thus, for a given Delay I cell *i, I*_*i,stim*_ = 0, *stim* ∈ {*targ, dist*}, throughout trials, while for a given Delay E cell *i*:

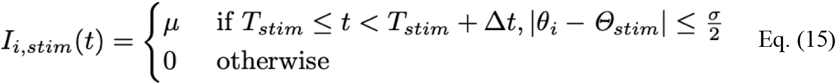

*Based on the analysis of the inverted-U response across stimulus strengths for Networks 1 and 2 in Figure S4; used in Figure 5 and for the analysis of the inverted-U response across distractor positions for Networks 1 and 2 in Figure S5.

**Based on the analysis of the inverted-U response across stimulus strengths for Network 3 in Figure S4, used for analysis of the inverted-U response across distractor positions for Network 3 in Figure S5, and for the I_*NEAR*_-only and I_*OPP*_-only networks in Figure S6.

#### Simulation protocol for visual fixation model

We also simulated three-second trials of a visual fixation task mimicking Mitchell *et al*.’s^**1**^ and Nandy *et al*.’s^**2**^ protocols, again without explicit saccadic responses. For a given trial, at a time point of *T*_*fix*_ = 400 ms, until the end of the trial, we applied a fixation stimulus as a current injection of strength *μ* = 0.20 nA onto each E^*FIX*^ cell. To test the distractor resistance of the microcircuit, we then applied 175 distractor stimuli starting at *T*_*dist*_ = 600 ms, also until the end of the trial, onto E^*CU*E^ cells. Specifically, each distractor stimulus was applied onto the respective E^*CU*E^ cells within arcs with widths of *σ* = 30° centred at angles *Θ*_*dist*_ that were randomly sampled from nine positions on the ring (0°, 45°, 90°, 135°, 180°, 225°, 270°, 315°). Each distractor stimulus had a duration of Δ*t* = 16 ms and was of the same strength as the fixation stimulus. The fixation with distractors period was 2400 ms until the end of the trial. Thus, for a given I^*CU*E^ or I^*FIX*^ cell *i, I*_*i,stim*_ = 0, *stim* ∈ {*fix, dist*}, throughout trials. For a given E^*CU*E^ cell *i, I*_*i,stim*_ is determined by Eq. 15. Notably, since the fixation stimulus has no angular position and is applied to all E^*FIX*^ cells, which themself lack angular position selectivity, for a given E^*FIX*^ cell *i*:

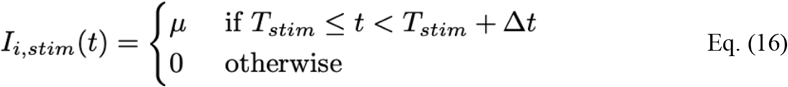

#### Classification protocol for visuospatial working memory model

To classify the relevant dynamical regimes of the networks in the visuospatial working memory model, representing stages of the inverted-U, as outcomes of task trials, we estimated the population vector^**49**, **50**^ from the activity of the Delay E cell population during three periods: (1) baseline, from 0.2 to 0.4 secs of a trial (i.e. following the initial 0.2 secs after trial onset, to allow activity to equilibrate into a steady state, and until target stimulus presentation), (2) target delay, from 2.2 to 2.7 secs (i.e., until distractor stimulus presentation), and (3) distractor delay, from 4.5 to 5.0 secs (i.e., until the end of the trial). Thus, if [*n*_*i*_, *i* = 1 … 2048] are the spike counts of all E cells, labelled by an angle [*θ*_*i*_, *i* = 1 … 2048], during the respective trial period, the population vector is the normalized sum of each E cell’s selectivity vector, 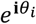, weighted by its spike count: 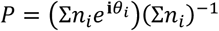, where the **i** in the exponential function is the imaginary unit. We extracted the population vector angle *θ*_act_ and modulus 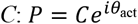. For each trial period, we took *θ*_act_ as the decoded angle. Given that *C* quantifies the signal-to-noise ratio, we took it as a measure of confidence in the decoded angle. Like Cano-Colino *et al*.^**51**^, we arbitrarily took a threshold of *C* > 0.5 as a criterion for confident decoding of *θ*_act_.

We classified four dynamical regimes in the visuospatial working memory model, characterized by (1) transient activity, (2) noise- and distractor-resistant persistent activity, (3) distractible persistent activity, and (4) erroneous activity using the population vector angle *θ*_act_ and modulus *C*. In the transient activity regime, the confidence of the decoded *θ*_act_ during either or both the target and the distractor delay period was low (*C* < 0.5). Confident decoding of *θ*_act_ in only one of the delay periods also indicated a transient activity regime. Notably, we ensured that the transient activity regime included only trials where the mean population firing activity of active Delay E cells (i.e. that spiked at least once) averaged over each period was ≤10 Hz. In the noise- and distractor-resistant persistent activity regime, we confidently decoded *θ*_act_ during both the target and distractor delay periods within +/-22.5° of the target position (0°). Notably, we excluded potential trials in which we may have confidently decoded *θ*_act_ within +/-22.5° of the target position during the baseline period (i.e. noise-evoked activity) due to uncertainty about whether any subsequent confident decoding of *θ*_act_ at the target position during the target and distractor delay periods is driven by the target stimulus or the noise-evoked activity. In the distractible persistent activity regime, we confidently decoded *θ*_act_ within +/-22.5° of the target position during the target delay period, but then also within +/-22.5° of the distractor position during the distractor delay period. Again, we excluded potential trials where we may have confidently decoded *θ*_act_ within +/-22.5° of the target position during the baseline period. The erroneous persistent activity regime accounted for any activity pattern that did not meet the conditions for the prior three regimes. For example, due to the disinhibition, at sub-optimal D1R occupancies across networks, errors mainly involved noise-evoked or spatially diffuse persistent activity, reflected in confidently decoded *θ*_act_ at non-target positions (> +/-22.5° of the target). Conversely, due to the overinhibition, at supra-optimal D1R occupancies across networks, errors mainly involved stimulus-evoked persistent activity but only for the target or for the distractor, reflected in confidently decoded *θ*_act_ at one but not both stimulus-specific positions (< +/-22.5°) during the respective period. Additionally, errors also occurred at the optimal D1R occupancy across networks, primarily when distractors appeared 67.5°-90° away from the target. These errors stemmed from the merging (vector summation) of the persistent activity traces for the target and the distractor due to their proximity during the distractor delay period, and the peak of the resulting activity trace taking a non-target position (> +/-22.5° of the target) intermediate to where the two external stimuli had appeared.

#### Classification protocol for visual fixation model

To classify the relevant dynamical regimes in the visual fixation model, we estimated the mean firing activity of the E^*FIX*^ cell population during three periods: (1) baseline, from -0.2 to 0 secs before trial onset (i.e. following the initial 0.2 secs after simulation onset (−0.4 to -0.2 secs before a trial), to allow activity to equilibrate into a steady state, and until fixation stimulus presentation), (2) fixation without distractors, from 0.4 to 0.6 secs (i.e., until distractor stimuli presentation), and (3) fixation with distractors, from 0.6 to 3.0 secs (i.e., until the end of the trial). Thus, if [*n*_*i*_, *i* = 1 … 256] are the spike counts of all E^*FIX*^ cells during the respective trial period (binned into 100 equally spaced intervals within each period), Δ*t* is the duration of a period, and *N* is the number of the E^*FIX*^ cells, the mean firing activity, 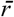, is the total sum of the spike counts, weighted by the duration of the given period and the number of E^*FIX*^ cells: 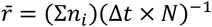.

We classified three dynamical regimes in the visual fixation model, characterized by (1) noise- and distractor-resistant persistent activity, (2) distractible persistent activity, and (3) erroneous activity, using the mean firing activity. Notably, classifying a transient activity regime in the E^*FIX*^ population is not meaningful because once turned on, the fixation stimulus remains on until the end of the trial. Thus, while underexcitation and overinhibition resulted in transient activity in the visuospatial working memory model (at no and very high D1R occupancy respectively), here in the visual fixation model these conditions would result in distractible persistent activity instead. In the noise- and distractor-resistant persistent activity regime, the mean population firing activity of all E^*FIX*^ cells averaged over the baseline period was ≤10 Hz, but >10 Hz over the fixation-without-distractors and fixation-with-distractors periods, and without any dips in the time-varying mean population firing rate ≤10 Hz in the period from 150 ms after the onset of distractors (to minimize the aftereffects of the fixation-without-distractors period) until the end of the trial. In the distractible persistent activity regime, the mean population firing activity of E^*FIX*^ cells was ≤10 Hz averaged over the baseline period, >10 Hz over the fixation-without-distractors-period, but the time-varying mean population firing rate dipped to ≤10 Hz at some point following the initial 150 ms after the onset of distractors and before the end of the trial. In the erroneous persistent activity regime, the mean population firing activity of E^*FIX*^ cells averaged over the baseline period was >10 Hz. Importantly, these regimes captured all trial outcomes.

Notably, characterizing the extreme regimes of the inverted-U by transient activity of the E^*FIX*^ cells is not meaningful here because overt sustained attention by definition requires the fixation stimulus to be on, whereas during working memory maintenance target stimuli are off. Thus, E^*FIX*^ cells could not be transiently active, unless there is extreme inhibition caused by D1R stimulation that is outside of the functionally feasible range of D1R occupancy on 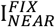 cells used here. Thus, any suboptimal D1R occupancy on 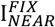 cells would only result in erroneous noise-evoked persistent activity in E^*FIX*^ cells due to insufficient feedback inhibition. Optimal D1R occupancy would ensure sufficient feedback inhibition to stabilize the persistent activity of E^*FIX*^ cells, as well as sufficient feedforward inhibition driven by the E^*CU*E^ cells. The latter would allow microcircuits to detect potentially relevant environmental stimuli that are significantly strong. Finally, any supraoptimal D1R occupancy would disrupt the persistent activity of E^*FIX*^ cells, primarily due to feedforward overinhibition driven by E^*CU*E^ cells that are activated by distractors, but also feedback overinhibition in the absence of distractors and given high enough D1R occupancy on 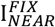 cells.

#### Analyses

For the analysis of the inverted-U response across stimulus strengths (Figure S4), we simulated 1000 trials for each unique parameter set, specifically for each network configuration (3), main D1R occupancy level (5), and stimulus strength (17, from 0 to 0.40 pA, in steps of 0.025). We applied the classification protocol to all trials and then grouped them into 1000 sets, each containing one trial from each of the five main D1R levels. For each network configuration, we then calculated the fraction of trial sets at each stimulus strength, in which the network reproduced the inverted-U response defined as (1) transient activity at no D1R occupancy, (2) any one of erroneous, noise- and distractor-resistant, or distractible persistent activity at low D1R occupancy, (3) noise- and distractor-resistant persistent activity at mid D1R occupancy, (4) distractible persistent activity at high D1R occupancy, and (5) transient activity at very high D1R occupancy.

For the analysis of the inverted-U response across distractor positions (Figure S5), we also simulated 1000 trials for each unique parameter set, specifically for each network configuration (3), main D1R occupancy level (5), and distractor position (7, from 45.0° to 180.0° in steps of 22.5°). Again, we applied the classification protocol to all trials. Additionally, we conducted the same analysis and classification on two alternative networks (Figure S6), which comprised only, respectively, Delay I_*NEAR*_ or Delay I_*OPP*_ cells, in addition to the Delay E cells. Importantly, in these alternative networks, we maintained the total number of 512 I cells to match the net inhibitory drive across the I_*NEAR*_-only and I_*OPP*_-only networks for comparison with Network 3. Moreover, since, necessarily, all I cells in each alterative network expressed D1Rs because reproducing the inverted-U response requires dopaminergic modulation of inhibition, we used the D1R modulation parameters from Network 3. To reiterate, studying any other more complex D1R expression patterns, here specifically within the I cell types, was prohibited by the combinatorial explosion of possible patterns.

For the granulated analysis of the effect of the species difference in D1R expression on PV neurons on visuospatial working memory maintenance in Network 1 (Figure 6), we again simulated 1000 trials for each unique parameter set. This included one case for the macaque-D1R model and three cases for the marmoset-D1R model, across 21 levels of D1R occupancy (from 0 to 1, in steps of 0.05). Once again, we applied the classification protocol to all trials.

For the granulated analysis of the effect of the species difference in D1R expression on PV neurons on visual fixation performance (Figure 8A1-2), we similarly simulated 1000 trials for each unique parameter set and species-D1R model, across the 21 levels of D1R occupancy on 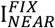 cells (from 0 to 1, in steps of 0.05). Again, we applied the classification protocol to all trials. Next, we calculated the mean fixation duration from the non-erroneous trials (i.e., excluding trials with erroneous outcomes) for each D1R occupancy level and species-D1R model (Figure 8B1). Then, we pooled all trials with non-erroneous outcomes from the five D1R occupancy levels forming the mid D1R occupancy range for each species-D1R model and computed the species-D1R model means (Figure 8B2). To evaluate whether the observed differences in the means could arise by chance, we performed a random permutation test with 10,000 iterations. Specifically, for each permutation, we concatenated the two species-D1R datasets for the mid D1R occupancy range, randomly permuted the combined data, split it back into two groups of the original sizes, and computed the difference in means. The p-value was determined as the fraction of permuted differences whose absolute value exceeded or equaled that of the observed difference in means.

Notably, within each analysis, the seed that generated the noise was different for each simulated trial, serving to avoid bias and simulate the natural variability inherent in biological systems. Specifically, we utilized the simulation ID number as the seed (see Python scripts), thereby also ensuring reproducibility.

#### Numerical integration

We used a second-order Runge-Kutta integration algorithm with a time step of Δ*t* = 0.1 ms for each simulation, which reportedly provided an optimal trade-off between computational tractability and precision in a similar spiking neural network^**52**^.

#### Code availability

The custom scripts for the modeling were written in Python 3.10 using the Brian2 (Version 2.5.1) neural simulator^**53**^ and are openly available on GitHub (https://github.com/tsvet-ivanov/marm-mac-D1R). Simulations that ran on the high-performance computing facilities used Python 3.7, 3.11, or 3.12.

## Acknowledgements

This research was funded by NSF 2015276 to AFTA, JMT, FMK and SAM, as well as by a Bristol Neuroscience of Mental Health award (co-funded by Steve Scobie and Anne Graham) and BBSRC BB/X013243/1 to SFW. We thank the Yale Center for Cellular & Molecular Imaging Confocal Facility for usage of their Zeiss LSM 880 Airyscan microscope. The modeling work was carried out using the high-performance computing facilities of the Advanced Computing Research Centre, University of Bristol – http://www.bristol.ac.uk/acrc/. We are grateful to Kristy Gibbs, Lisa Ciavarella, Tracy Sadlon, Sam Johnson, and Michelle Wilson for their technical expertise. Additionally, we thank Albert Compte, Joao Barbosa, Paul Anastasiades, John Hongyu Meng, Christos Constantinidis, Susheel Vijayraghavan, Loïc Magrou, Shengtao Yang, and Min Wang for helpful discussions, as well as Matt Jones, Ulysse Klatzmann, Aswathi Thrivikraman, Dabal Pedamonti, and Eva Sevenster for feedback on the manuscript. Further thanks to Benjamin Corrigan and Diego Buitrago-Piza for their behavioral task illustrations, which we have adapted for Figure 1.

## Notes

### Competing Interest Statement

The authors have declared no competing interest.

### Summary of Updates

Addition of new modeling results, including new main figures, main text, methods, and supplemental figures

