## Supplemental Figures for "Dopamine D1 receptor expression in prefrontal parvalbumin neurons influences distractibility across species"

### Supplementary Materials

#### Supplementary Figure 1

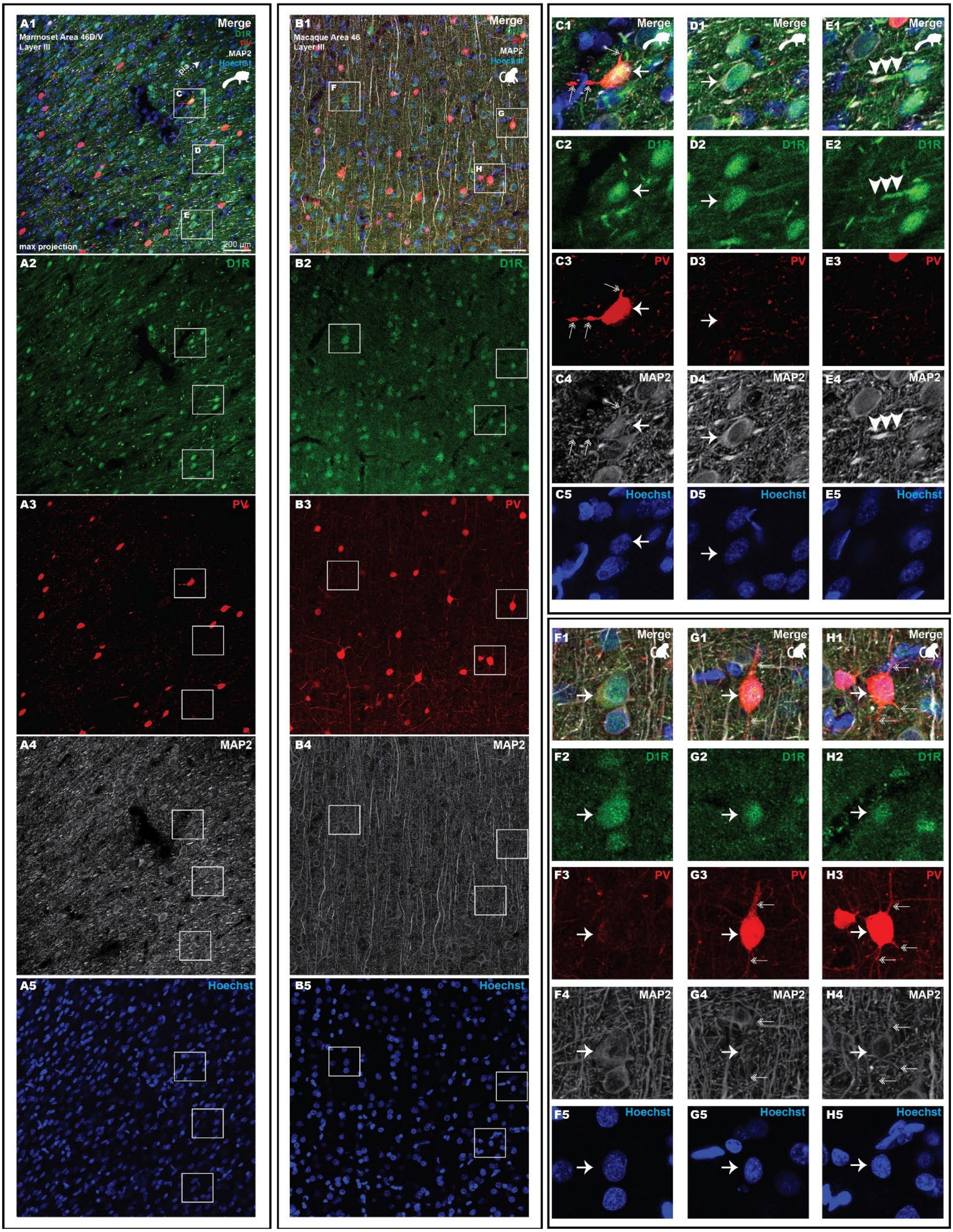

**Figure S1 – Comparative multi-label immunofluorescence in layer III dlPFC for PV, D1R, MAP2, and Hoechst.** Marmoset **(A1-5)** and macaque **(B1-5)** layer III dlPFC ***A1-A5,*** Layer III of marmoset A46D, labeled for PV (red), D1R (green), microtubule-associated protein 2 (MAP2, white), and the Hoechst nuclear stain (blue). White square insets are shown on the right. Apical dendrites were cut in oblique cross-section, distinguishable using MAP2 labeling **(A4)**, and are oriented toward Layer I in the direction of the upper right. Apical dendrites are also strongly labeled by the D1R antibody **(A2)**, and overlap with the MAP2 label (**E1**, **E2**, **E4**, white arrowheads). Macaque apical dendrites are oriented upward **(B1)**, and are strongly labeled by MAP2 **(B4)**, although their apical dendrites are labeled more weakly for D1R than in the marmoset **(B2)**. In general, PV neuronal processes were more prominently labeled in the macaque compared to the marmoset (**B3** vs. **B4**). ***C1-C5,*** A marmoset D1R positive PV neuron (white arrow). The PV soma and proximal PV processes (double-headed arrows) are also lightly labeled by MAP2 **(C4)**. Strong D1R labeling corresponds with the nucleus (**C2** vs. **C5**). ***D1-D5,*** A D1R positive pyramidal neuron (**D2**, white arrow) with an apical dendrite, stretching toward the left, in the marmoset. The soma and apical dendrite are also labeled strongly by MAP2 **(D4)**. Strongest D1R labeling was observed in the nucleus (**D2** vs. **D5**). ***E1-E5,*** A strongly labeled D1R segment (white arrowheads), which was common in the marmoset tissue, likely an obliquely cut apical dendrite. The D1R label **(E2)** overlaps and is continuous with the MAP2 label **(E4)**. ***F1-F5,*** A D1R positive pyramidal neuron **(F2)** in the macaque, with an apical dendritic segment reaching upwards toward Layer I. The neuron is also labeled by MAP2 **(F4)**. D1R labeling is strongest in the nuclear region (**F2** vs. **F5**). ***G1-G5,*** A macaque PV neuron **(G3)** with robust D1R labeling **(G2)**. The D1R labeling is strongest in what corresponds to the nuclear region (**G2** vs. **G5**). The PV neuron is negative for MAP2 **(G4)**. ***H1-H5,*** A macaque PV neuron **(H3)** lightly positive for D1R (white arrow). To its left is a PV neuron negative for D1R. The center PV neuron is positive for MAP2 **(H4)**, including proximal PV processes (double-headed arrows in **H2**, **H4**). Again, the D1R is strongest in the nuclear region (**H2** vs. **H5**).

**Supplementary Figure 2**

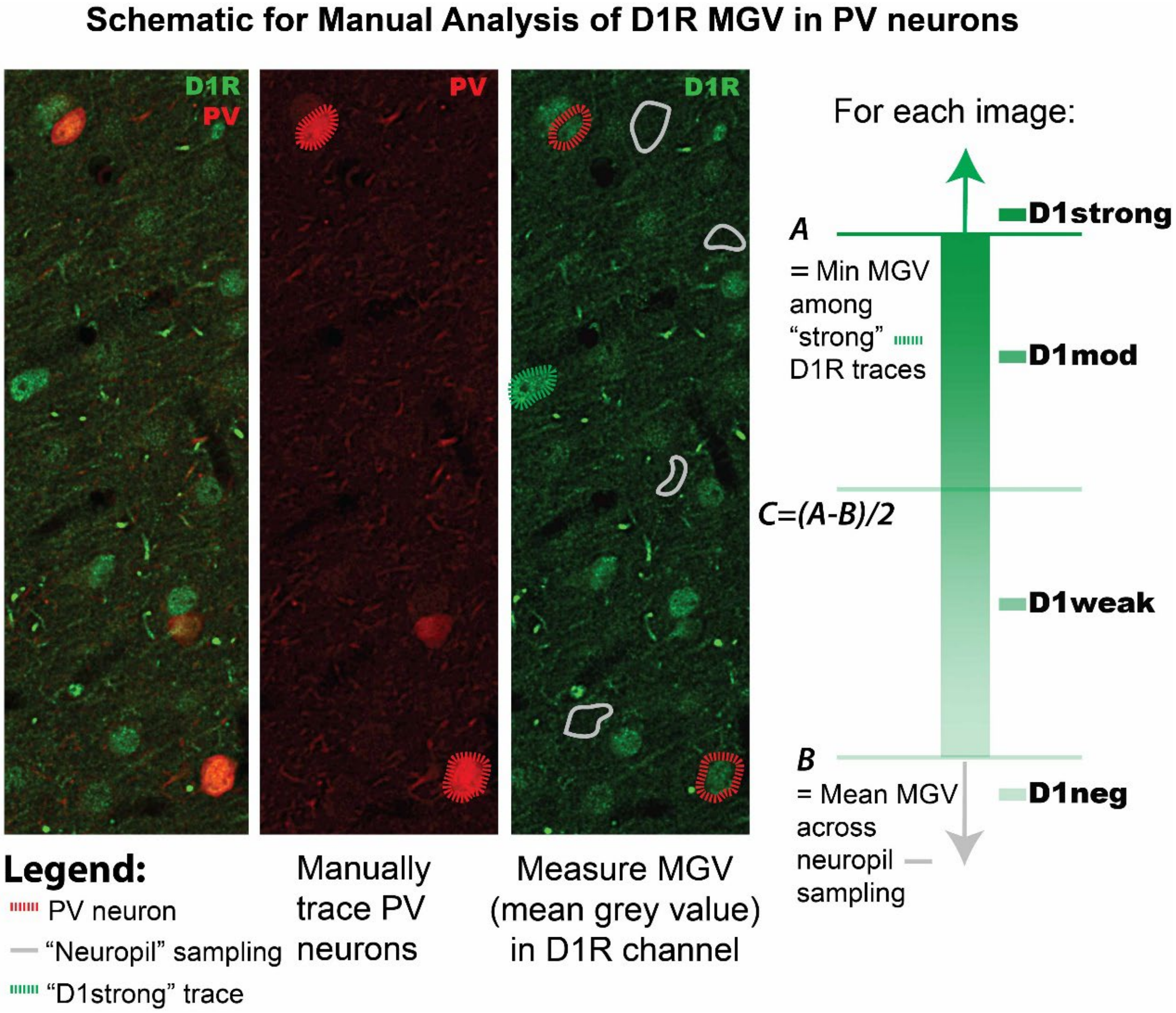


**Figure S2 – Simplified exemplary schematic for manual comparative analysis of D1R expression levels in PV neurons.** Single focal planes from z-stacks were extracted (left image). The PV channel was isolated (middle column, red), and the somata of PV neurons were traced (red dashed line). PV neurons not completely in plane were discarded (e.g., faint PV+ neuron in the middle of the image). In the isolated D1R channel (green, right image), strongly labeled D1R neurons were traced (dashed green line), as well as regions of "neuropil"-like immunonegative labeling (grey outlines). The mean grey value (MGV) of the D1R signal was measured in each type of trace. A binning strategy was used to classify PV traces (schematic at right). PV cells were deemed "D1neg" if their MGV fell below the average MGV from the sampled neuropil traces. PV cells were deemed "D1strong" if their average MGV exceeded the minimum from strongly labeled D1R cells in the image. PV cells in the intermediate bins were classified based on their average MGV falling between those two extremes. This analysis was performed for each image to ensure normalization based on the unique illumination level for each plane. Data obtained per image was then compiled across images by subject.

**Supplementary Figure 3**

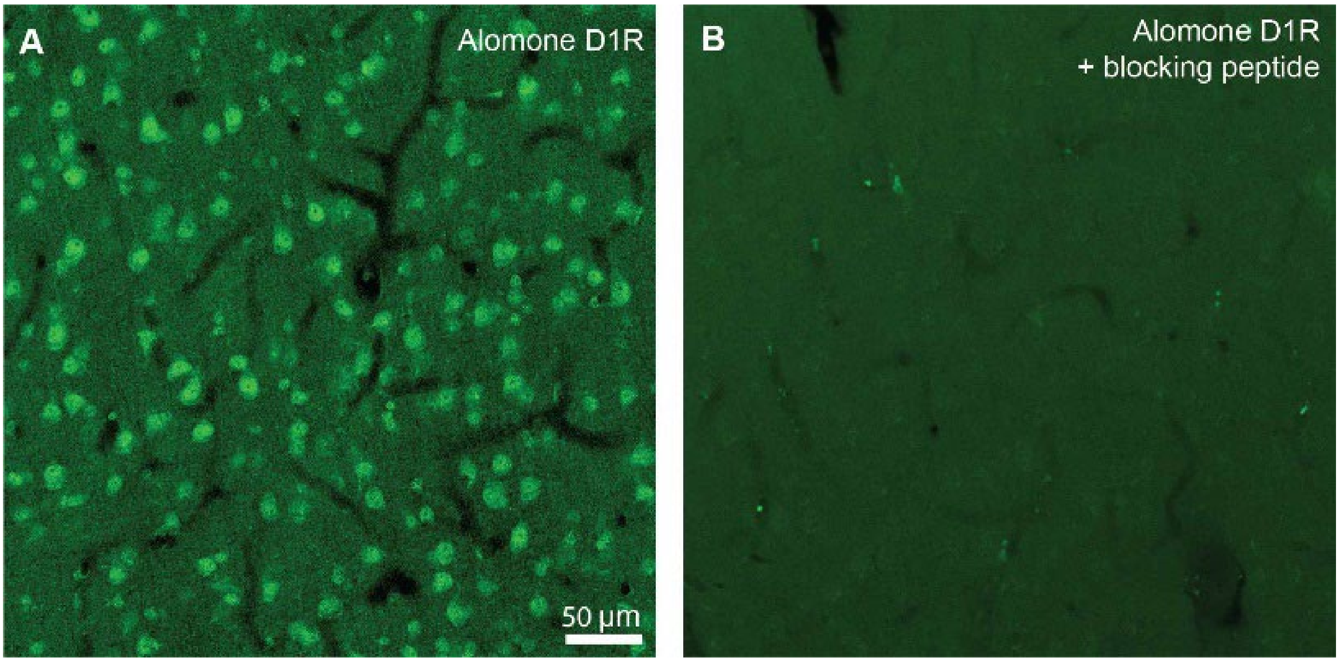


**Figure S3 - Preadsorption Control for D1R antibody.** ***A,*** Macaque dlPFC labeled with Alomone rabbit anti-D1R (cat# ADR-001). Strong labeling is visible particularly in pyramidal-like cells with apical dendrites oriented upwards toward Layer I. ***B,*** Test section using preadsorption control for primary antibody. Co-incubation of the primary antibody with the bespoke blocking peptide (Alomone, #BLP-DR001) produced negligible labeling.

**Supplementary Figure 4**

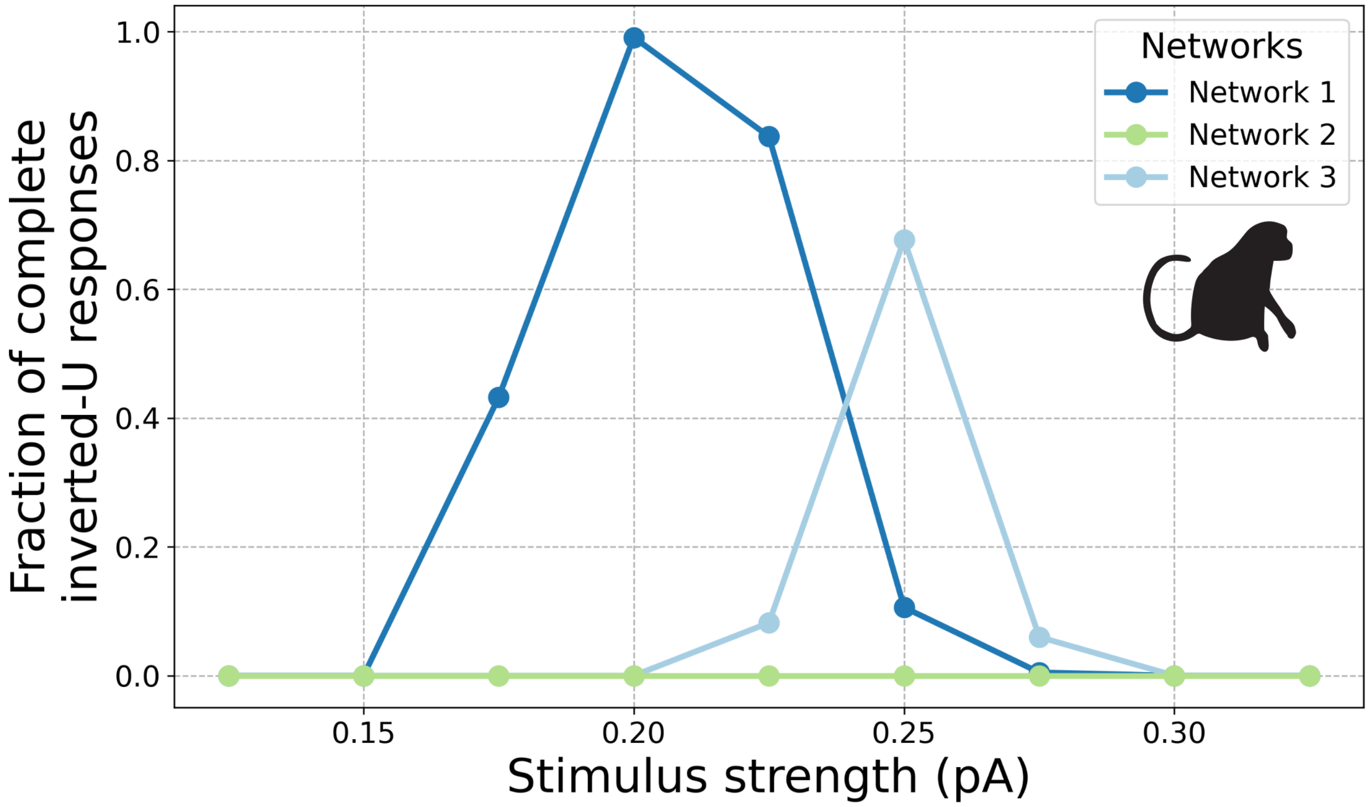


**Figure S4 – D1R expression in Delay E and $I_{NEAR}$ cells facilitates the inverted-U response**. The plot presents the fraction of complete inverted-U responses in the macaque-D1R configuration of the visuospatial working memory model, out of 1000 repetitions for each combination of cell type-specific D1R expression pattern (Networks 1-3), stimulus strength, and set of five main D1R occupancy levels (no, low, mid, high, very high). A complete inverted-U involves (1) transient activity at no D1R occupancy, (2) any one of erroneous, noise- and distractor-resistant, or distractible persistent activity at low D1R occupancy, (3) noise- and distractor-resistant persistent activity at mid D1R occupancy, (4) distractible persistent activity at high D1R occupancy, and (5) transient activity at very high D1R occupancy. Network 1 could reproduce the inverted-U for approximately the 0.150-0.300 pA range of stimulus strengths, with a frequency of 0% for 0.150 pA, 43.2% for 0.175 pA, 99.1% for 0.200 pA, 83.7% for 0.225 pA, 10.6% for 0.250 pA, 0.5% for 0.275%, and 0% for 0.300 pA. Network 2 could not reproduce the inverted-U response at any stimulus strength. Network 3 could reproduce the inverted-U for approximately the 0.200-0.300 pA range of stimulus strengths, with a frequency of 0.0% for 0.200 pA, 8.2% for 0.225 pA, 67.7% for 0.250 pA, 6.0% for 0.275 pA, and 0% for 0.300 pA.

**Supplementary Figure 5**

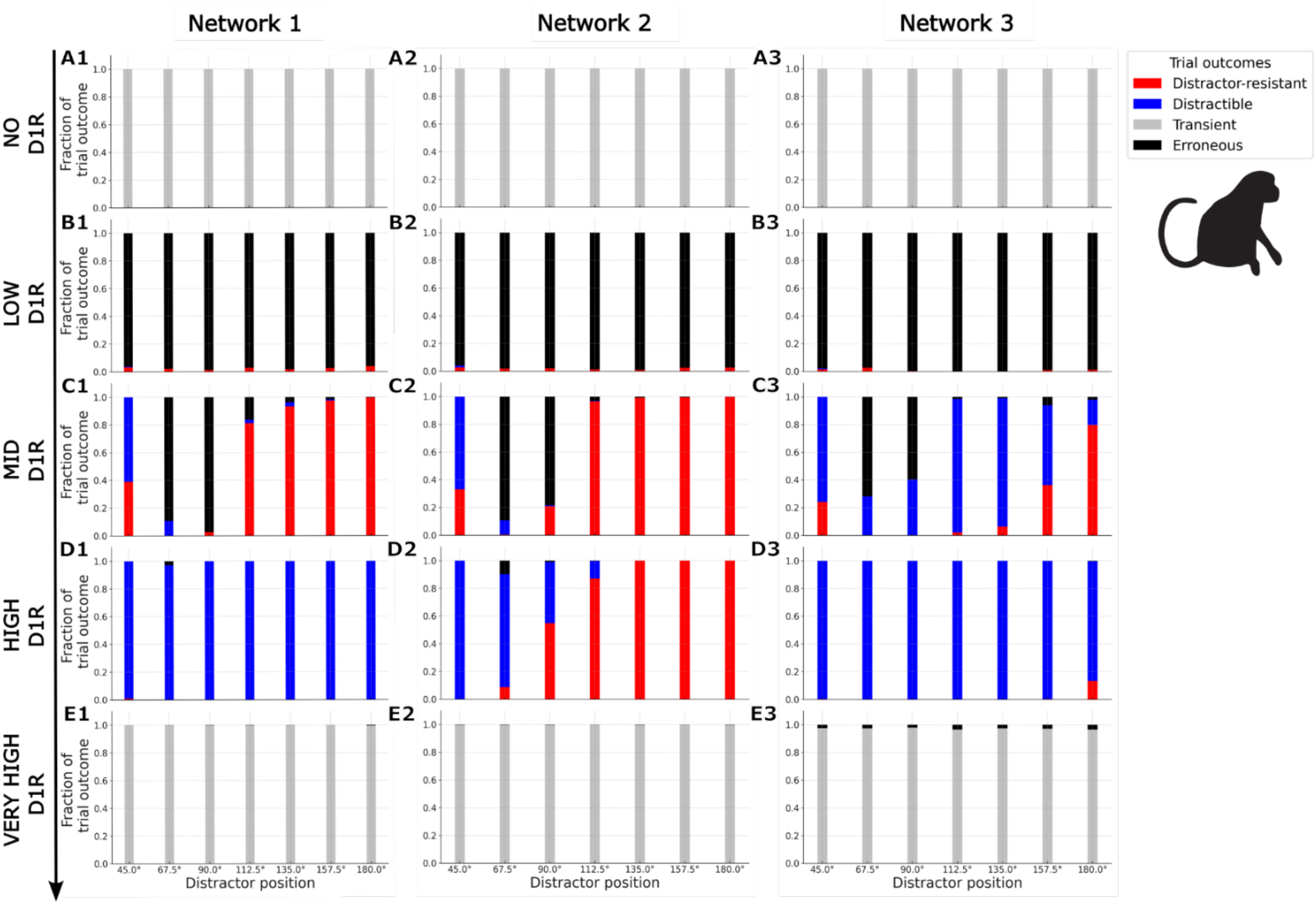


**Figure S5 – Fraction of classified outcomes 1000 of trials at different distractor positions across the main D1R occupancy levels for Networks 1, 2, and 3 in the macaque-D1R configuration. The strength of the stimuli is 0.2 pA for Network 1 and 2, and 0.25 pA for Network 3.** ***A,*** At no D1R occupancy, Networks 1, 2, and 3 **(A1-3)** exhibit transient activity across distractor positions. ***B,*** At low D1R occupancy, all three networks **(B1-3)** overwhelmingly exhibit erroneous persistent activity across distractor positions, with some steady fraction of noise- and distractor-resistant persistent activity. ***C,*** At mid D1R occupancy, Networks 1 and 2 **(C1-2)** alike exhibit predominantly distractible persistent activity, but also a significant fraction of noise- and distractor-resistant persistent activity, at near distractor positions (45.0°). For both networks, erroneous persistent activity predominates at near-to-intermediate positions (67.5°-90.0°), with some decreasing fraction at intermediate-to-far positions (112.5°-180.0°). Both networks overwhelmingly exhibit noise- and distractor-resistant persistent activity at intermediate-to-far positions (112.5°-180.0°). Conversely, Network 3 **(C3)** predominantly exhibits distractible persistent activity at near (45.0°) and intermediate-to-far (112.5°-157.5°) positions, erroneous persistent activity at near-to-intermediate positions (67.5°-90.0°), and noise- and distractor-resistant persistent activity only at the farthest position (180.0°). Upon inspection of several individual example trials, the erroneous persistent activity at near-to-intermediate positions (67.5°-90.0°) across the networks appears to result from the persistent activity peak shifting to a position between the target and distractor during the distractor delay period. ***D,*** At high D1R occupancy, Networks 1 and 3 **(D1, D3)** overwhelmingly exhibit distractible persistent activity across distractor positions. Conversely, Network 2 **(D2)** exhibits overwhelmingly distractible persistent activity at near distractor positions (45.0°-67.5°), gradually switching to a robust noise- and distractor-resistant persistent activity regime at intermediate-to-far distractor positions (90.0°-180.0°). ***E,*** At very high D1R occupancy, all three networks **(E1-3)** overwhelmingly exhibit transient activity across distractor positions, with some fraction of erroneous persistent activity, especially in Network 3.

**Supplementary Figure 6**

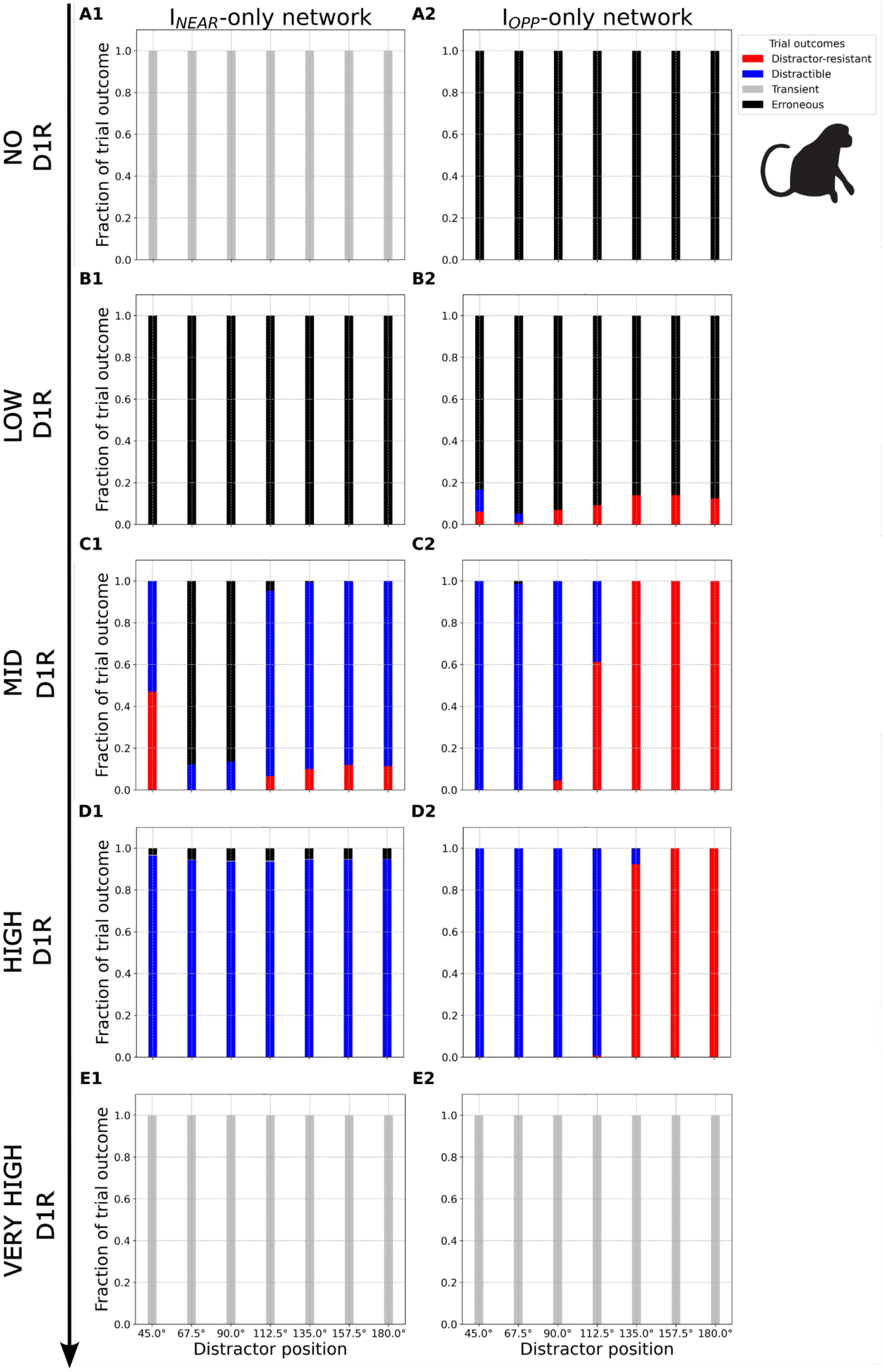

**Figure S6 – Fraction of classified outcomes of 1000 trials at different distractor positions across the main D1R occupancy levels for the $I_{NEAR}$-only and the $I_{OPP}$-only networks in the macaque-D1R configuration. The strength of the stimuli is 0.25 pA.** ***A,*** At no D1R occupancy, the $I_{NEAR}$-only network **(A1)** exhibits transient activity across distractor positions. The $I_{OPP}$-only network **(A2)** exhibits erroneous persistent activity across distractor positions. Upon inspection of several individual example trials, the errors mainly consisted of stimulus-evoked persistent activity but only for the target or for the distractor, similar to the overinhibition errors at higher D1R occupancies in Networks 1-3. ***B,*** At low D1R occupancy, the $I_{NEAR}$-only network **(B1)** exhibits erroneous persistent activity across distractor positions. The $I_{OPP}$-only network **(B2)** also primarily exhibits erroneous persistent activity across distractor positions, but also some fractions of distractible persistent activity at near positions (45.0°-67.5°), as well as noise- and distractor-resistant persistent activity with fractions across distractor positions forming a discernible irregular amplitude sinusoidal pattern. ***C,*** At mid D1R occupancy, the $I_{NEAR}$-only network **(C1)** exhibits distractible persistent activity primarily at near (45.0°) and intermediate-to-far (112.5°-180°), but also some at near-to-intermediate (67.5°-90.0°), distractor positions. There is a significant fraction of noise- and distractor-resistant persistent activity at the near (45.0°) and some at the intermediate-to-far (112.5°-180°) positions. Notably, erroneous persistent activity predominates at near-to-intermediate positions (67.5°-90.0°), with some gradually decreasing fractions at intermediate-to-far positions (112.5°-180.0°). Qualitatively different, the $I_{OPP}$-only network **(C2)** overwhelmingly exhibits distractible persistent activity at near-to-intermediate positions (45.0°-90.0°), switching to a robust noise- and distractor-resistant persistent activity regime at intermediate-to-far positions (112.5°-180.0°). ***D,*** At high D1R occupancy, the $I_{NEAR}$-only network **(D1)** overwhelmingly exhibits distractible persistent activity, with some steady fraction of erroneous persistent activity, across distractor positions. Conversely, the $I_{OPP}$-only network **(D2)** exhibits overwhelmingly distractible persistent activity at near-to-intermediate positions (45.0°-112.5°), abruptly switching to a robust noise- and distractor-resistant persistent activity regime at intermediate-to-far positions (135.0°-180.0°). ***E,*** At very high D1R occupancy, both networks **(E1-2)** exhibit transient activity across distractor positions. Importantly, focusing on the typical, mid D1R occupancy, $I_{NEAR}$-only inhibition facilitates significant resistance to near distractor stimuli and noise-evoked activations (45°), while $I_{OPP}$-only inhibition – strong resistance to far distractor stimuli and noise-evoked activations (112.5°-180.0°). $I_{OPP}$-only inhibition also curbs errors at the near-to-intermediate distractor positions (67.5°-90.0°). However, a microcircuit with only $I_{NEAR}$ or $I_{OPP}$ cells seems dysfunctional due to overall poor distractor resistance across distractor positions expected at mid D1R occupancy in the former case and a lack of distractibility for far distractors expected at high D1R occupancy in the latter case.

**Supplementary Figure 7**

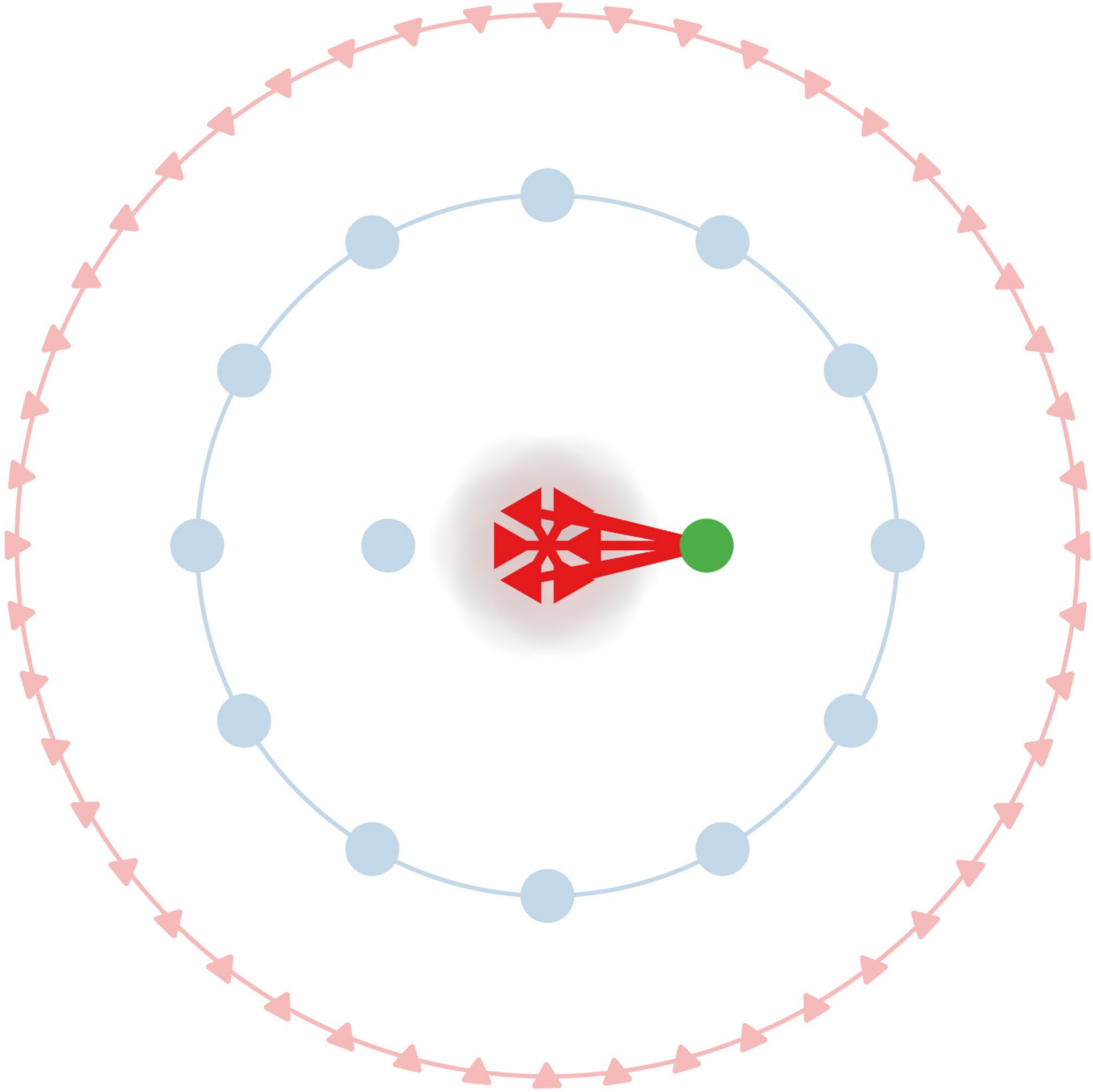

**Figure S7 – Simplified schematic representation of the Fixation Rule E cells driving an example Fixation $I_{OPP}$ cell.**
